# Highly ordered clustering of TNFα and BAFF ligand-receptor-adaptor complexes bound to lipid membranes

**DOI:** 10.1101/2024.07.22.604697

**Authors:** Chan Seok Lim, Jisun Lee, Ji Won Kim, Jie-Oh Lee

## Abstract

We report the cryo-EM structures of clusters of TNF receptor family proteins, TNFR1 and BAFFR. The receptor-ligand complexes were anchored to a flat lipid layer to mimic the membrane-bound state. We observed that the TNFα-TNFR1 complex forms highly ordered binary, bent, trigonal, and linear quadruple clusters of trimers on the lipid membrane. A non-competitive antagonist of TNFR1 disrupted these clusters without interfering with ligand binding. Moreover, we found that the BAFF-BAFFR complex forms pentagonal, double-pentagonal, or half-spherical clusters of trimers. Mutations in BAFF that inhibit BAFFR receptor activation prevented ordered clustering without disrupting receptor binding. TRAF3 induced a structural shift in the BAFF-BAFFR cluster, resulting in a flat hexagonal cluster. Our data demonstrated that precisely structured clustering is essential for the activation of these receptors. The lipid monolayer method will aid in studying the clusters of other transmembrane proteins and facilitate the discovery of therapeutic agents that regulate their clustering.

## INTRODUCTION

The tumor necrosis factor (TNF) family of proteins plays several critical roles in regulating immune responses, inducing apoptosis, regulating cell proliferation and differentiation, and maintaining tissue homeostasis ^1^. Humans possess 19 TNF family ligands and 29 TNF receptor family members ^2,3^. The binding of TNF family ligands to their specific receptors leads to the activation of transcription factors, such as NF-κB and AP-1, which can regulate gene expression and promote various cellular responses. TNF family ligands and receptors are among the most crucial targets for drug discovery ^4^. For example, antibodies against TNFα, including adalimumab, named Humira commercially, are being used as anti-inflammatory drugs ^5^.

TNFα is the prototypical member of the TNF ligand family that can bind to two receptors, namely TNFR1 and TNFR2 ^6^. The structures of TNFα bound to the extracellular domain of TNFR1 have been elucidated using X-ray crystallography ^7,8^. In the crystal structure, TNFα is a trimer arranged as a triangular cone, with each molecule in contact with the other two. The extracellular domain of TNFR1 is an elongated molecule composed of four disulfide-containing motifs, known as cysteine-rich repeats (CRDs), each comprising approximately 40 amino acids ^9^. The TNFα trimer binds to three receptor molecules, one at each of the three TNF monomer-monomer interfaces. The binding of the trimeric ligand induces receptor trimerization and activates downstream signaling pathways by recruiting adaptor proteins, including TRADD ^10,11^. The binding of TNFα to TNFR2 recruits TRAF2 instead of TRADD and activates the NF-κB and JNK pathways ^12,13^.

BAFF is a member of the TNF-ligand family. It promotes the survival and maturation of B cells and regulates the selection of the B cell repertoire ^14–16^. It is typically elevated in autoimmune diseases and serves as a target for the treatment of systemic lupus erythematosus ^17^. Soluble BAFF exists in two forms. Similar to other members of the TNF family, BAFF forms a trimeric complex in solution. Under certain conditions, BAFF trimers oligomerize to form a cage-like structure comprising 60 subunits ^18,19^. A loop region, termed “flap”, facilitates the formation of the BAFF cage ^18–21^. This oligomerization increases the potency of BAFF in promoting B cell survival and maturation. BAFF has three receptors, namely BAFFR, TACI, and BCMA, which contain one or two CRDs in their extracellular domains ^22–24^. Activated BAFFR recruits TRAF3, which subsequently negatively regulates NF-κB function ^25–27^.

TRAF proteins play essential roles in the intracellular signal transduction of several receptor families, including TNF, IL-1. Toll-like and NOD-like receptors ^28^. Upon receptor activation, TRAFs are directly or indirectly recruited to the intracellular domains of the receptors. They subsequently engage other signaling proteins, which ultimately activate transcription factors such as NF-κB and AP-1 to induce immune and inflammatory responses and confer protection from apoptosis. TRAF family proteins exhibit a modular organizational structure characteristic of adaptor proteins that function as docking assembly links for structurally dissimilar factors ^29,30^. The C-terminal half of TRAF binds to receptors when activated. It can be further divided into two sections: the coiled-coil and TRAF-C domains. The coiled-coil domain folds into a parallel and triple coiled-coil structure, which enables the trimerization of TRAF proteins. The TRAF-C domain directly binds to the intracellular domains of the receptor ^31–33^. The N-terminal half of TRAF proteins is more divergent, although all TRAF proteins except TRAF1 feature five closely spaced Zinc-finger domains. The N-terminal regions of TRAF2 and 6 dimerize both in solution as well as their crystal structures ^34,35^. It has been proposed that they cross-link the trimeric C-terminal regions, inducing the formation of an extensive hexagonal network of TRAF proteins ^36^.

Previous studies proposed that the trimeric TNFα-TNFR1 and BAFF-BAFFR complexes form higher-order aggregates ^1,18,19,37–39^. However, high-resolution structures of these aggregates have not yet been reported, partly because the membrane-bound state of the receptor complexes could not be reproduced for X-ray crystallographic or cryo-electron microscopy (cryo-EM) studies using truncated or detergent-solubilized receptors. To overcome this technical hurdle, we adopted a lipid monolayer method to mimic the membrane-anchored state of the proteins ^40^. Using this method, we could determine the clustered structures of the TNFα-TNFR1 and BAFF-BAFFR complexes. We showed that the trimeric TNFα-TNFR1 units form several highly ordered clusters with binary, bent, trigonal, and linear quadruple shapes. Trimeric BAFF-BAFFR complexes were assembled into ordered pentagonal, double-pentagonal, and half-spherical clusters on the membrane. We also showed that the binding of TRAF3 rearranges the BAFF-BAFFR cluster into a flat hexagonal cluster.

## RESULTS

### Lipid monolayer method

Membrane receptors must be attached to lipid membranes to understand their ligand-binding and activation mechanisms fully. We adopted the lipid monolayer method, as reported by Truong *et al.* and others, for this purpose ^40^. The lipid monolayer method has long been used to generate two-dimensional protein crystals for electron crystallographic studies or concentrate samples on EM grids for affinity grid preparation ^41^. In this method, the hydrophobic fatty acids in the lipid layer point to the air and the hydrophilic head groups are aligned to face the aqueous solution. We used a phospholipid monolayer containing 5%–20% 18:1 DGS-NTA(Ni^2+^) phospholipids to mimic the membrane-bound state of the receptors (Figure S1A). The lipid monolayer was subsequently incubated with receptor complexes tagged with hexa- or octa-histidine sequences. The binding of the histidine tag to the Ni-NTA head group mediates the anchoring of the receptor complexes to the lipid layer (Figure 1A). The lipid layer confines the diffusion of receptor complexes within the viscous and two-dimensional membranes, which enhances the lateral interaction between the receptor complexes. Compared to other artificial lipid layers mimicking cellular membranes, such as liposomes, bicelles, or nanodiscs ^42^, the lipid monolayer has an advantage in generating a large and flat area of lipid membranes. After protein binding, the lipid monolayer was transferred onto a cryo-EM grid and imaged (Figure S1A).

**Figure 1.**
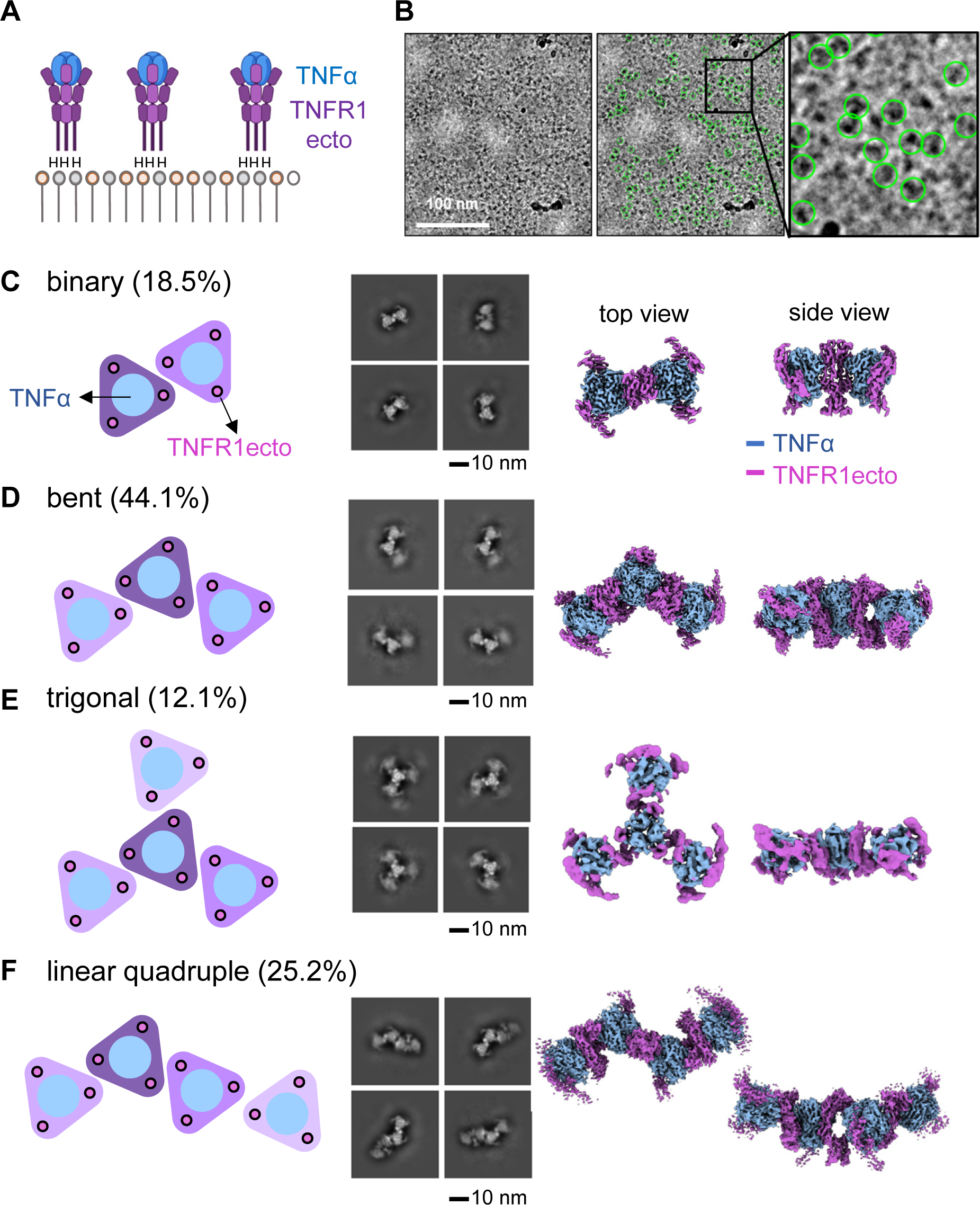
The TNF*α*-TNFR1ecto complex forms ordered clusters on the lipid layer. (A) An octa-histidine tag was attached to the C-terminus of TNFR1ecto. Following this, the TNFα-TNFR1ecto complex was bound to the Ni-NTA lipid monolayer. (B) A representative cryo-electron microscopy (cryo-EM) image of the TNFα-TNFR1ecto complex. The protein particles used for structure determination are marked with green circles. (C) A schematic diagram (left), 2D class averages (middle), and 3D refined map (right) of the binary cluster. The ratio of the protein particles that belong to the binary cluster is written in parenthesis. A schematic diagram, 2D class averages, and 3D refined map of the (D) bent cluster, (E) trigonal cluster, and (F) linear quadruple cluster

### Clustering of the TNF*α*-TNFR1 ectodomain complex on the lipid monolayer

To determine the structure of the TNFα-TNFR1 complex in the membrane-bound state, an octa-histidine tag was fused to the C-terminus of the ectodomain of TNFR1, termed TNFR1ecto (Figures 1A, S1B and Table S1). Subsequently, the TNFα-TNFR1ecto complex was generated by mixing TNFR1ecto with soluble TNFα. After purification, the receptor-ligand complex was bound to the Ni-NTA lipid monolayer, as shown in Figure 1A, and the structure was determined by cryo-EM (Table S2). Cryo-EM images of the complex revealed a clustered pattern (Figure 1B). The 2D class averaged images of the protein particles showed four major classes of structures. In the binary cluster, dimerized TNFR1 receptors bridged two trimeric TNFα-TNFR1ecto complexes, forming a linear structure containing six ligands bound to receptors in a 1:1 molar ratio (Figure 1C). This binary cluster served as the basic unit for the formation of larger clusters. In the bent cluster, two TNFR1 receptors in the central trimer formed dimeric interactions with TNFR1 receptors from the neighboring TNFα-TNFR1 trimeric units (Figure 1D). In the trigonal cluster, three binary clusters were assembled around a central TNFα-TNFR1ecto trimer (Figure 1E). A linear quadruple cluster was formed by connecting two binary clusters (Figure 1F). Among these various clusters, the bent cluster, containing nine ligands and nine receptors, was the most abundant, comprising 44.1% of the total protein clusters.

Dimerization between two TNFR1 receptors in neighboring trimeric units mediated the formation of all clusters (Figure 2A). The TNFα ligands are not directly involved in cluster formation. The two TNFα-TNFR1ecto trimers in the binary cluster were rotated by 63.7° when viewed from the side. The TNFR1 dimer interface structures were identical in all cluster forms and could be divided into two areas. The first dimer interface, named the “PLAD” region in previous literature, was primarily formed by CRD1 of TNFR1 (Figure S2A) ^43^. The network of ionic bonds formed by K19, H34, K35, D49, and E64 plays a major role at this CRD1 interface. Mutation of K19 or K32 to alanine has been demonstrated to disrupt receptor dimerization ^39,43^. K19 is directly involved in receptor dimerization in the structure. Conversely, K32 plays a role in stabilizing the CRD1 structure by interacting with E64 and is, therefore, indirectly involved in receptor dimerization. The second interface was formed by CRD4 of TNFR1. The ionic interactions between E131 and K132 and the hydrophobic interactions between L127 and V136 play key roles at this dimerization interface.

**Figure 2.**
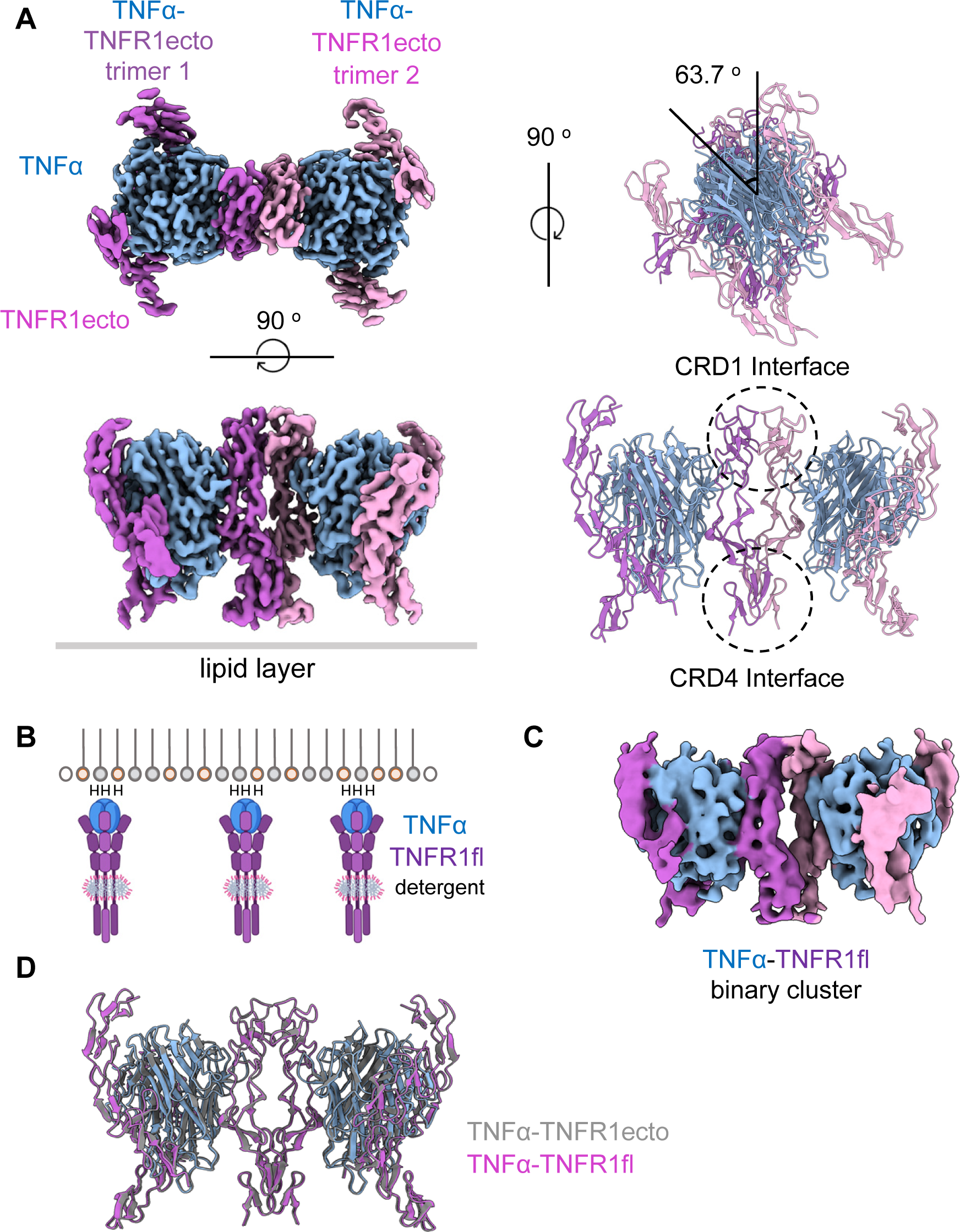
Structures of the binary clusters of the TNF*α*-TNFR1ecto and the TNF*α*-TNFR1fl complexes. (A) Cryo-EM electron density map and the overall structure of the binary cluster of the TNFα-TNFR1ecto complex. (B) The tethering of the TNFα-TNFR1fl complex to the Ni-NTA lipid monolayer. A hexa-histidine tag was attached to the C-terminus of TNFα. (C) Cryo-EM electron density map of the binary cluster of the TNFα-TNFR1fl complex. (D) Structural comparison of the binary clusters of the TNFα-TNFR1ecto and TNFα-TNFR1fl complexes.

The dimerization of TNFR1 was previously reported using X-ray crystallographic analysis ^38^. They determined the structure of the TNFR1 ectodomain in the absence of bound ligands. This dimeric receptor structure is an artifact of crystallization that cannot be reproduced in solution. However, notably, the dimeric structure of TNFR1ecto in the crystal state can be superimposed onto that of its clustered state on the lipid membrane, showing a Cα r.m.s.d. of 1.7 Å (Figure S2B).

### Clustering of the full-length TNFR1-TNF*α* complex on the lipid membrane

We produced a full-length TNFR1 receptor, named TNFR1fl (Figure S1B and Table S1), to evaluate whether the transmembrane or intracellular domain of TNFR1 is involved in the clustering of the TNFα-TNFR1 complex and determined the TNFα-TNFR1fl complex structure (Table S2). We fused a hexa-histidine tag to the N-terminus of the ligand and bound the receptor-ligand complex to the Ni-NTA lipid monolayer (Figure 2B and Table S1). The 2D class averages of the cryo-EM images show that the TNFα-TNFR1fl complex primarily formed a binary cluster with a similar structure to that of the TNFα-TNFR1ecto complex (Figure S2C). The structure of the binary TNFα-TNFR1fl cluster was determined at a resolution of 6.0 Å (Table S2). The cryo-EM map corresponding to the transmembrane and intracellular domains was not clearly visible, presumably because of the structural flexibility of the connecting linker regions between the extracellular, transmembrane, and intracellular domains (Figures 2C and S2C). The binary cluster structures of the ectodomain and full-length TNFR1-TNFα complexes were practically identical, and the two structures could be superimposed with a Cα r.m.s.d. of 0.416 Å (Figure 2D). This observation highlights that the extracellular domain of the receptor plays a major role in clustering and that the intracellular and transmembrane domains have minimal contributions, if any, to cluster formation. Trigonal and linear quadruple clusters were not observed for the TNFα-TNFR1fl complex. This could be attributed to the small amount of detergent present in the buffer used to solubilize the full-length receptors. This detergent may have weakened the interactions between the receptors. Alternatively, this could be because the receptors are not directly attached to the lipid membrane and may have increased mobility, which destabilizes larger clusters (Figure 2B).

### Disruption of TNF*α*-TNFR1 clusters by a non-competitive antagonist

DOM1h-574-208 is a nanobody that has been demonstrated to inhibit the activation of the TNFR1 receptor without disrupting TNFα binding or receptor trimerization ^44^. Furthermore, it can significantly reduce IL-8 production in HEK293 cells transfected with TNFR1 ^45^. The long-lasting variant of DOM1h-574-208, named DMS5541 or TNFR1-AlbudAb, demonstrated effectiveness in KYM-1D4 cell cytotoxicity assays and dose-dependent VCAM-1 upregulation in HUVECs ^46^. Additionally, it also reduced osteoclast formation in *ex vivo* human rheumatoid synovial membrane cell cultures ^47^. However, the precise mechanism of action of this non-competitive antagonist has not yet been elucidated. We determined the structure of the TNFα-TNFR1ecto-DOM1h-574-208 complex in a solution and a lipid-bound state to understand the structural basis of DOM1h-574-208-mediated antagonism (Figure 3 and Table S2). The structure in a solution state revealed that the DOM1h nanobody was bound to the CRD4 region of TNFR1 (Figure 3A). The binding site of DOM1h is located opposite to that of the ligand-binding site. Therefore, nanobody did not disturb ligand binding or receptor trimerization. Nanobody binding weakened the TNFR1-TNFR1 dimeric interaction in the membrane-bound state, and approximately 60% of the receptor cluster dissolved into isolated TNFα-TNFR1ecto trimeric units. The remaining 40% of nanobody-bound proteins exhibited a binary-clustered structure (Figure 3B). The low-resolution cryo-EM map of this binary cluster clearly demonstrated that the nanobody disrupts the CRD4 interface of the TNFR1 dimer, and it was observed that two trimeric TNFα-TNFR1 units were rotated by approximately 60° compared to the binary cluster with no bound nanobody (Figure 3C). The cluster disruption was not due to structural changes in the receptor, as evidenced by the fact that the receptor structures with and without the bound nanobody could be superimposed with a Cα r.m.s.d. of 0.684 Å (Figure 3D). Our structural observation and reported antagonistic activity of the DOM1h nanobody demonstrate that proper clustering is indispensable for the activation of TNFR1 by TNFα.

**Figure 3.**
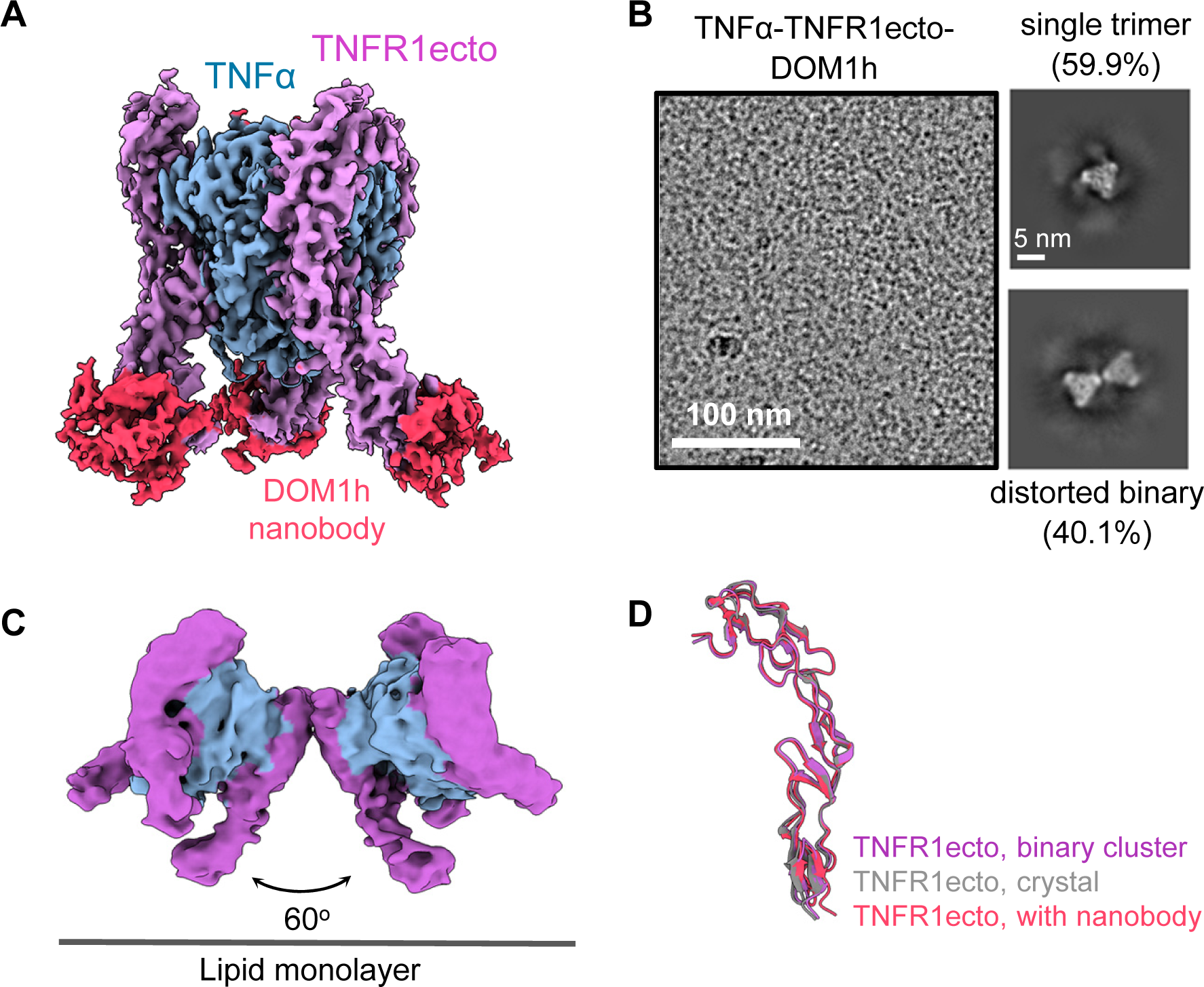
DOM1h-574-208 nanobody distorts the structure of the binary cluster of the TNF*α*-TNFR1ecto complex. (A) A composite electron density map of the TNFα-TNFR1ecto-DOM1h nanobody complex in a solution state. Focused refinement was performed on one of the three DOM1h nanobody regions of the map. Subsequently, maps for the remaining two nanobodies were generated by applying threefold symmetry. (B) A representative image (left) and 2D class averages (right) of the TNFα-TNFR1ecto-DOM1h nanobody complex bound to the lipid monolayer. (C) The cryo-EM electron density map of the binary cluster of the TNFα-TNFR1ecto-DOM1h nanobody complex bound to the Ni-NTA lipid layer. The position of the bound nanobodies is not visible in this low-resolution map, presumably owing to structural flexibility in the CRD4 region of the TNFR1 receptor. (D) Superimposition of the TNFR1ecto structures.

### Clustering of the BAFF-BAFFR complex on the lipid monolayer

An octa-histidine tag was fused to the C-terminus of the BAFFR ectodomain, termed BAFFRecto, to determine the structure of the BAFF-BAFFR complex in the membrane-bound state (Figure S1B and Table S1). The BAFF-BAFFRecto complex was formed by mixing BAFFRecto with soluble BAFF. After purification, the receptor-ligand complex was bound to the Ni-NTA lipid monolayer, as shown in Figure 4A. The structure of the complex was determined by cryo-EM (Table S2). The BAFF-BAFFRecto complex, which is trimeric in solution (Figure S3A), mainly formed three types of clusters when attached to the lipid monolayer (Figure 4A, lower panels). The dominant form was a pentagonal ring composed of five trimeric units of BAFF-BAFFRecto. However, a significant number of double pentagons and half spheres were also formed. We did not observe any particles with the complete globular cage form of the BAFF-BAFFRecto complex. As a control, the purified full-sized cage form of BAFF was attached to BAFFRecto bound to the lipid monolayer (Figure S3B). Approximately half of the BAFF cage was dissolved into the pentagonal cluster (Figure S3C). This finding suggests that the full-sized BAFF cage becomes unstable and transforms into a pentagonal cluster when bound to BAFFR embedded in the lipid membrane.

**Figure 4.**
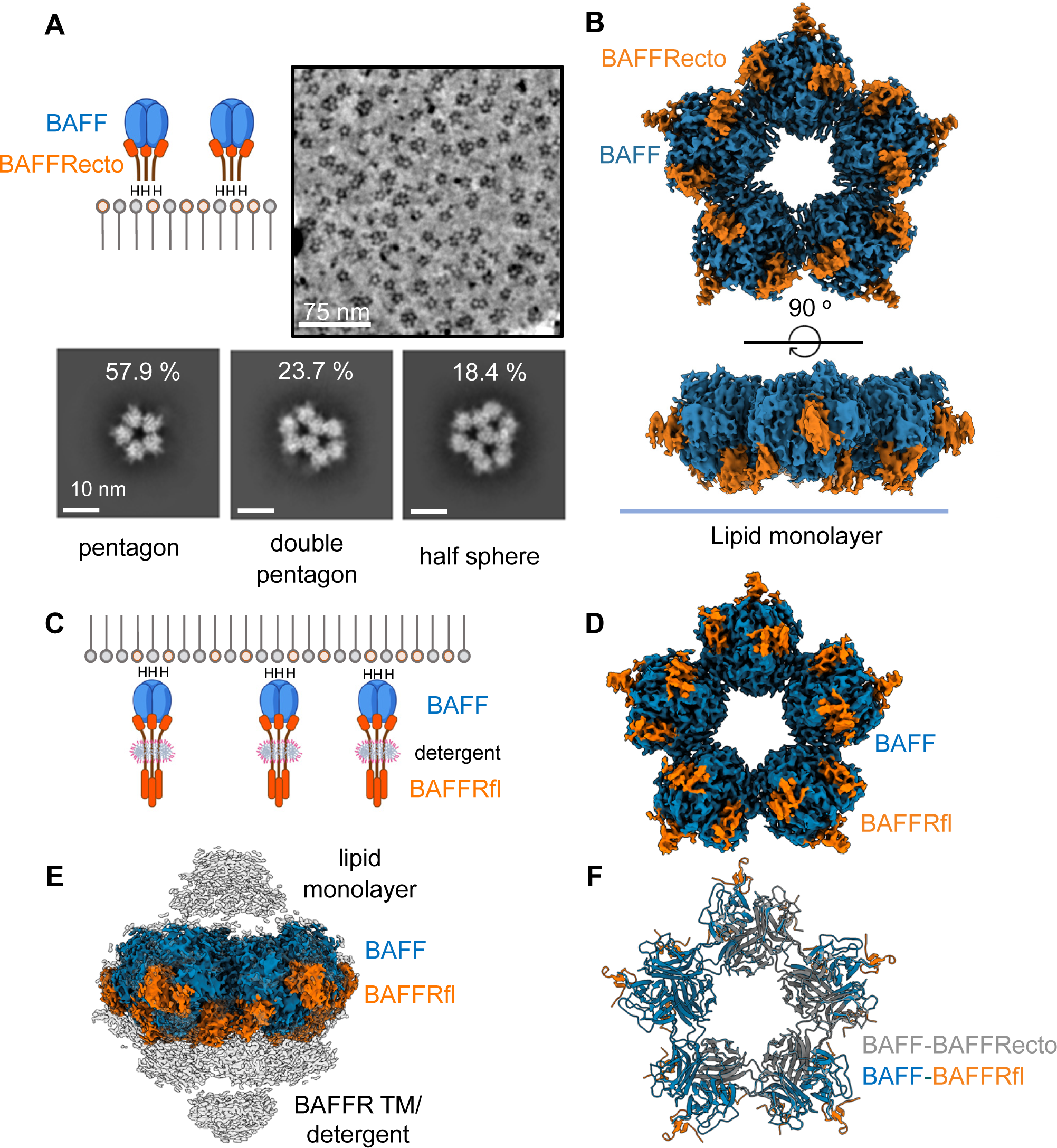
The BAFF-BAFFRecto complex forms an ordered cluster on the lipid layer. (A) An octa-histidine tag was attached to the C-terminus of the BAFFRecto. Following this, the BAFF-BAFFRecto complex was bound to the Ni-NTA lipid monolayer. A representative cryo-EM image (upper) and the 2D class averages (lower three panels) of the clusters are shown. (B) The cryo-EM electron density map of the pentagonal cluster. (C) A hexa-histidine tag was attached to the N-terminus of BAFF. Subsequently, the BAFF-BAFFRfl complex was bound to the Ni-NTA lipid monolayer. (D) The cryo-EM map of the pentagonal BAFF-BAFFRfl cluster. (E) Cryo-EM maps of BAFF-BAFFRfl. The lower and higher contour level maps are colored in grey and blue/orange, respectively. (F) Structural comparison of the pentagonal BAFF-BAFFRecto and BAFF-BAFFRfl clusters.

The pentagonal cluster structure of the BAFF-BAFFRecto complex was determined at a resolution of 3.3 Å (Figure 4B and Table S2). We found that the assembly of the pentagonal ring in the structure was primarily mediated by interactions between the two “flap” regions of the BAFF trimer (Figure S4A). Ionic interactions between R214, K216, D222, and E223 play a central role in this dimeric interaction. This region plays a key role in the formation of full-sized BAFF cages, as shown in previous crystallographic studies ^18–20^. We observed that the structure of the pentagonal cluster of BAFF-BAFFRecto was practically identical to the pentagonal substructure of the full-sized BAFF-BAFFRecto cage, and the two structures could be superimposed with a Cα r.m.s.d. of 1.17 Å (Figure S4B, left panel). A double pentagonal ring was formed by the fusion of two pentagonal clusters (Figure 4A). Additionally, the fusion of the three pentagons led to the formation of a half-spherical BAFF-BAFFRecto cluster. The double-pentagon and half-sphere cluster structures could be superimposed onto the full-sized cage structure, albeit with significantly higher Cα r.m.s.d. values because the outer parts of the cluster structures shifted downwards toward the lipid membrane by approximately 20 Å (Figure S4B, middle and right panels). It appears that the flattening of the cluster, which is necessary for the receptors to attach to the lipid membrane, was responsible for the structural differences.

### Clustering of the full-length BAFFR-BAFF complex on the lipid membrane

We produced a full-length BAFFR, named BAFFRfl, to evaluate whether the transmembrane or intracellular domain of BAFFR induces structural changes in the BAFF-BAFFRecto clusters (Tables S1 and S2). We fused a hexa-histidine tag to the N-terminus of BAFF and bound the receptor-ligand complex to the Ni-NTA lipid monolayer (Figure 4C). The 2D class averages of the cryo-EM images revealed that the BAFF-BAFFRfl complex primarily formed pentagonal clusters with structures similar to those of the BAFF-BAFFRecto complex (Figure S4C). The cryo-EM map corresponding to the transmembrane and intracellular domains of BAFF-BAFFRfl was not clearly visible, presumably because of the structural flexibility of the connecting linker regions between the extracellular, transmembrane and intracellular domains (Figure 4D and E). Structural differences between the pentagonal clusters of the ectodomain and full-length BAFF-BAFFR complexes were negligible, and the two structures could be superimposed with a Cα r.m.s.d. of 0.189 Å (Figure 4F). These data demonstrate that the extracellular domain of the receptor plays a major role in the clustering of the receptor-ligand complexes, whereas the intracellular or transmembrane domain has only minimal contributions, if any, to cluster formation.

### Disruption of BAFF-BAFFR clustering by mutations in the flap region

Previous crystallographic studies showed that the “flap” region in BAFF mediates the formation of the full-size cage in the crystal and solution states ^18–20^. Two mutations in this region, H242A and E247K, have been demonstrated to interfere with the biological activity of mouse BAFF without disrupting its binding to the BAFFR receptor ^21^. The study revealed that these mutations blocked BAFF signaling in reporter cell assays, disrupted B-cell maturation in the spleen, and altered the B/T cell ratio in the lymph nodes of knock-in mouse models. Among these, E247K exhibited the most significant inhibitory effect on BAFF function. H242 and E247 residues of mouse BAFF correspond to H218 and E223, respectively, in human BAFF. These mutations were introduced into the BAFF ligands to investigate whether they interfere with the formation of receptor-ligand clusters on the lipid layers and to determine the structure of the BAFF-BAFFRecto complex bound to the Ni-NTA lipid layer (Figure 5). The wild-type and mutant complex structures were determined using cryo-EM (Table S2). Conservative K216R and H218A mutations with milder inhibitory effects partially disrupted the pentagonal clusters. However, for the E223K mutation, which exhibited the most severe inhibitory effect in mice, the trimeric units still aggregated; however, the ordered structure of the cluster was completely disrupted, and, consequently, disordered aggregates were formed. These data demonstrate that receptor clustering with a properly ordered structure is essential for the activation of BAFF signaling.

**Figure 5.**
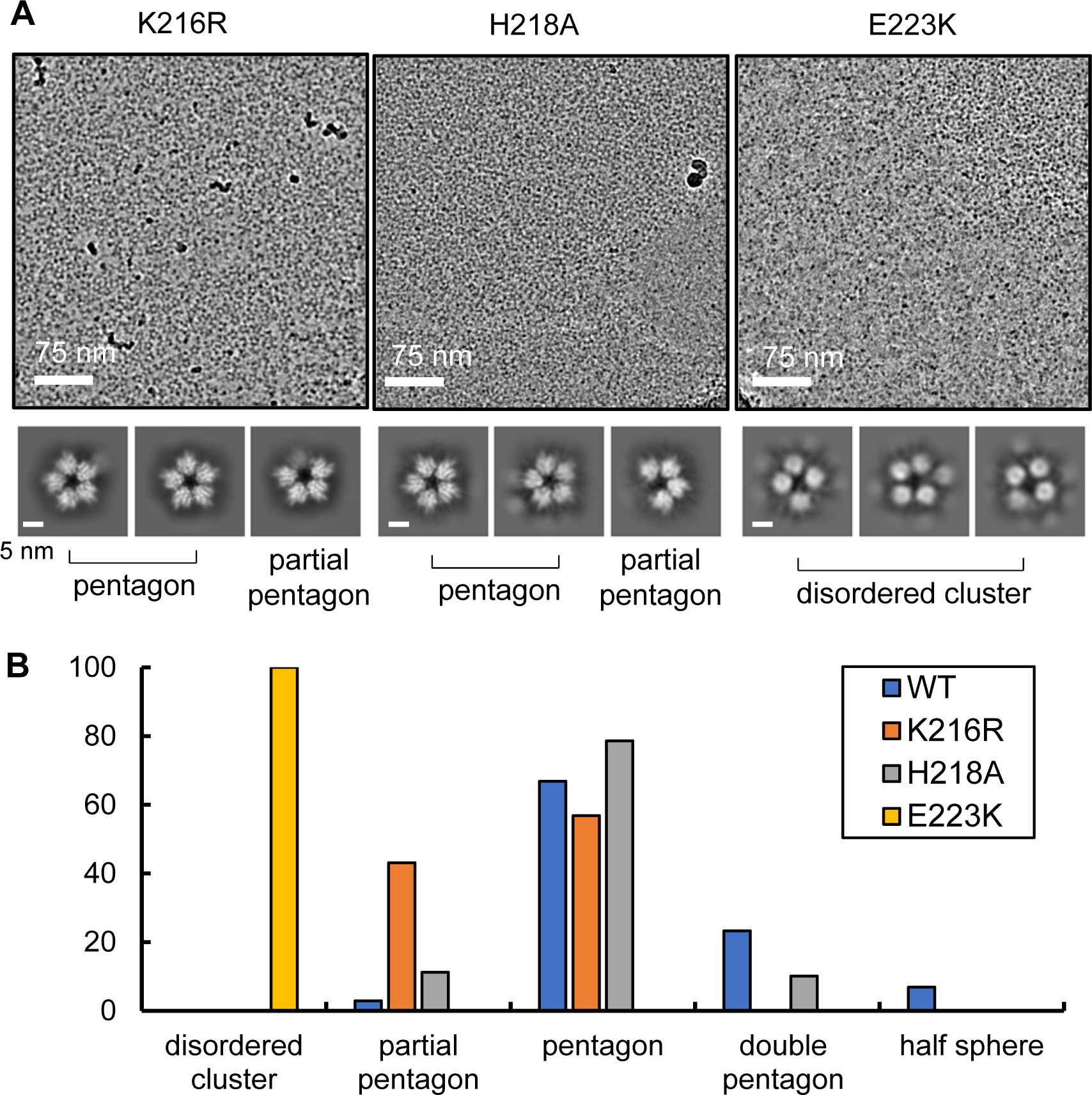
Mutations in the “flap” region of BAFF disrupt the cluster structures. (A) Representative cryo-EM images (upper) and 2D class averages (lower three panels) of the mutant BAFF-BAFFRecto complexes. (B) Clustering statistics of the BAFFRecto-BAFF mutant complexes.

### TRAF3 binding induces a structural shift in the BAFF-BAFFR cluster

We conducted a lipid monolayer experiment using soluble BAFF, full-length BAFFR, and truncated TRAF3 to explore the effects of TRAF3 binding to the BAFF-BAFFR cluster (Figure S1B and Table S1). The N-terminal RING and Zinc finger domains, which are dispensable for receptor binding, were removed from TRAF3. For membrane attachment, the BAFF ligand was tagged with a hexa-histidine sequence at the N-terminus and mixed with detergent-solubilized full-length BAFFR (Figure 6A). The histidine-tagged BAFF was named hisBAFF (Table S1). This mimicked the membrane-bound form of BAFF because full-length BAFF contains a transmembrane domain at the N-terminus of soluble BAFF (Figure S1B). The membrane-bound form of BAFF exhibits biological activity comparable to that of soluble BAFF ^22,23^. After forming the hisBAFF-BAFFRfl complex, truncated TRAF3 was added to the mixture, and the complex structure bound to the lipid monolayer was determined using cryo-EM (Figure 6A and Table S2).

**Figure 6.**
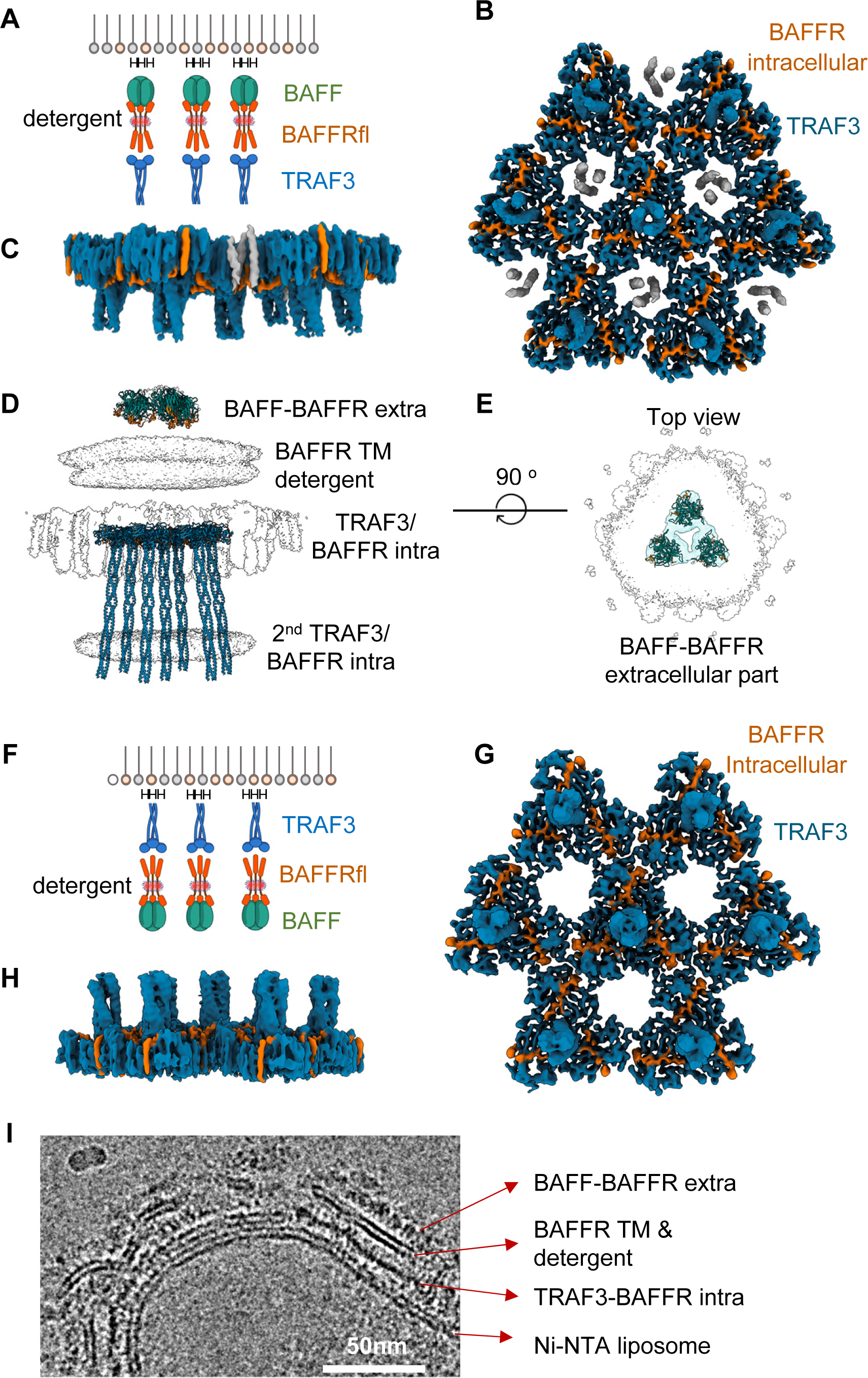
Cluster structures of the BAFF-BAFFRfl-TRAF3 complexes. (A) A hexa-histidine tag was attached to the N-terminus of BAFF. Subsequently, the hisBAFF-BAFFRfl-TRAF3 complex was bound to the Ni-NTA lipid monolayer. (B) Bottom view and (C) Side view of the cryo-EM map of the hisBAFF-BAFFRfl-TRAF3 cluster. Focused refinement of the TRAF3 cluster region was carried out. (D) Side view of the cryo-EM map of the hisBAFF-BAFFRfl-TRAF3 cluster without the focused refinement of the TRAF3 region. The structures of BAFF, BAFFR, and TRAF3 are fitted onto the map. Owing to structural flexibility, only part of the coiled-coil helices of TRAF3 are visible on the map. The missing parts of the coiled-coil helices are generated by AlphaFold 3. (E) Top view of the cryo-EM map of the hisBAFF-BAFFRfl-TRAF3 cluster without the focused refinement. The map corresponding to the extracellular part of the hisBAFF-BAFFR-TRAF3 cluster is shown in light green. The structures of the three BAFF-BAFFRecto trimers are fitted to the map. (F) An octa-histidine tag was attached to the N-terminus of TRAF3. Following this, the BAFF-BAFFRfl-hisTRAF3 complex was bound to the Ni-NTA lipid monolayer. (G) Top view and (H) Side view of the electron density map of the BAFF-BAFFRfl-hisTRAF3 cluster. (I) A representative cryo-EM image showing the side view of the BAFF-BAFFRfl-hisTRAF3 cluster. The image contains a large liposome that spontaneously formed during the preparation of the lipid monolayer.

Previous crystallographic studies have shown that the trimerized coiled-coil domain of TRAF3 protrudes as long rods. This coiled-coil rod served as a convenient landmark for analysis of the TRAF3 cluster because it appeared as a bright white dot in the cryo-EM 2D class averages (Figure S5A). As shown in Figure 6B and C, the hisBAFF-BAFFRfl-TRAF3 complex formed a large and flat hexagonal lattice. Owing to the structural flexibility of the linkers between extracellular, transmembrane, and intracellular domains of the receptor, only the short intracellular tail of BAFFR with 17 amino acids bound to TRAF3 was clearly visible in the map. The BAFFR intracellular domain structure within the cluster essentially resembled the crystal structure of the BAFF intracellular peptide bound to truncated TRAF3, and the two structures could be superimposed with a Cα r.m.s.d. of 1.27 Å (Figure S5B). The electron density map for the extracellular portion of the BAFF-BAFFR complex was only partially visible because of the structural flexibility of the linkers between the domains in the receptors. This low-resolution map also showed the hexagonal arrangement of the BAFF-BAFFR trimers (Figure 6D and E). Unexpectedly, we observed additional TRAF3 helical rods, depicted as gray densities in Figures 6B, C and S5C, intercalating the hexagonal lattice in an inverted orientation. This additional TRAF3 layer is likely an artifact resulting from the truncation of TRAF3 because, in full-length TRAF3, the N-terminal Zinc finger domains can prevent the intercalation of a second TRAF3 layer (Figure S1B).

We moved the histidine tag from the ligand to the N-terminus of truncated TRAF3 to further confirm the structural shift induced by TRAF3 binding (Figure 6F and Table S1) and determined its structure using cryo-EM (Table S2). We observed a hexagonal TRAF3 lattice in the cryo-EM images and 2D class averages (Figure S5D). The 3D reconstructed structure also showed a hexagonal TRAF3-BAFFR intracellular domain cluster (Figure 6G and H). Furthermore, no second TRAF3 layer was observed in this structure (Figure 6G, H, and I). The structure of this BAFF-BAFFRfl-hisTRAF3 cluster was nearly identical to that of hisBAFF-BAFFRfl-TRAF3 and the two structures could be superimposed with a Cα r.m.s.d. of 1.17 Å (Figure S5E). This structural observation confirms that hexagonal clustering is due to TRAF3 binding to BAFF-BAFFR and that the second layer of TRAF3 shown in Figures 6B and C had no effect on BAFF-BAFFR-TRAF3 cluster formation.

In BAFF-BAFFR-TRAF3 clustering, the TRAF-C and BAFFR intracellular domains play a central role. The coiled-coil domains of TRAF3 were separated by 76 Å in the cluster and did not contribute to cluster formation (Figure 6C and H). P164”, A165”, and T166” of BAFFR play a key role in TRAF3 clustering by simultaneously interacting with T469 and H470 in the first TRAF3 trimer and D463’, F474’, and P535’ in the second TRAF3 trimer (Figure 7A and B). Three M465 side chains of the TRAF3 trimers were assembled at the center of the TRAF3 cluster and formed hydrophobic interactions. In addition, we found that T541 and N545 in the first TRAF3 trimer and N517’, S518’, and S519’ in the second TRAF3 trimer formed a hydrogen bonding network and stabilized the TRAF3 cluster. Based on these structural observations, we propose a model for BAFFR receptor activation (Figure 7C). The binding of trimeric ligands induces receptor trimerization. Following this, the receptor-ligand trimers form mainly pentagonal clusters on the membrane. The binding of the intracellular adaptor TRAF3 to BAFFR induces a structural shift in the receptor cluster to a flat hexagonal lattice and initiates intracellular signaling.

**Figure 7.**
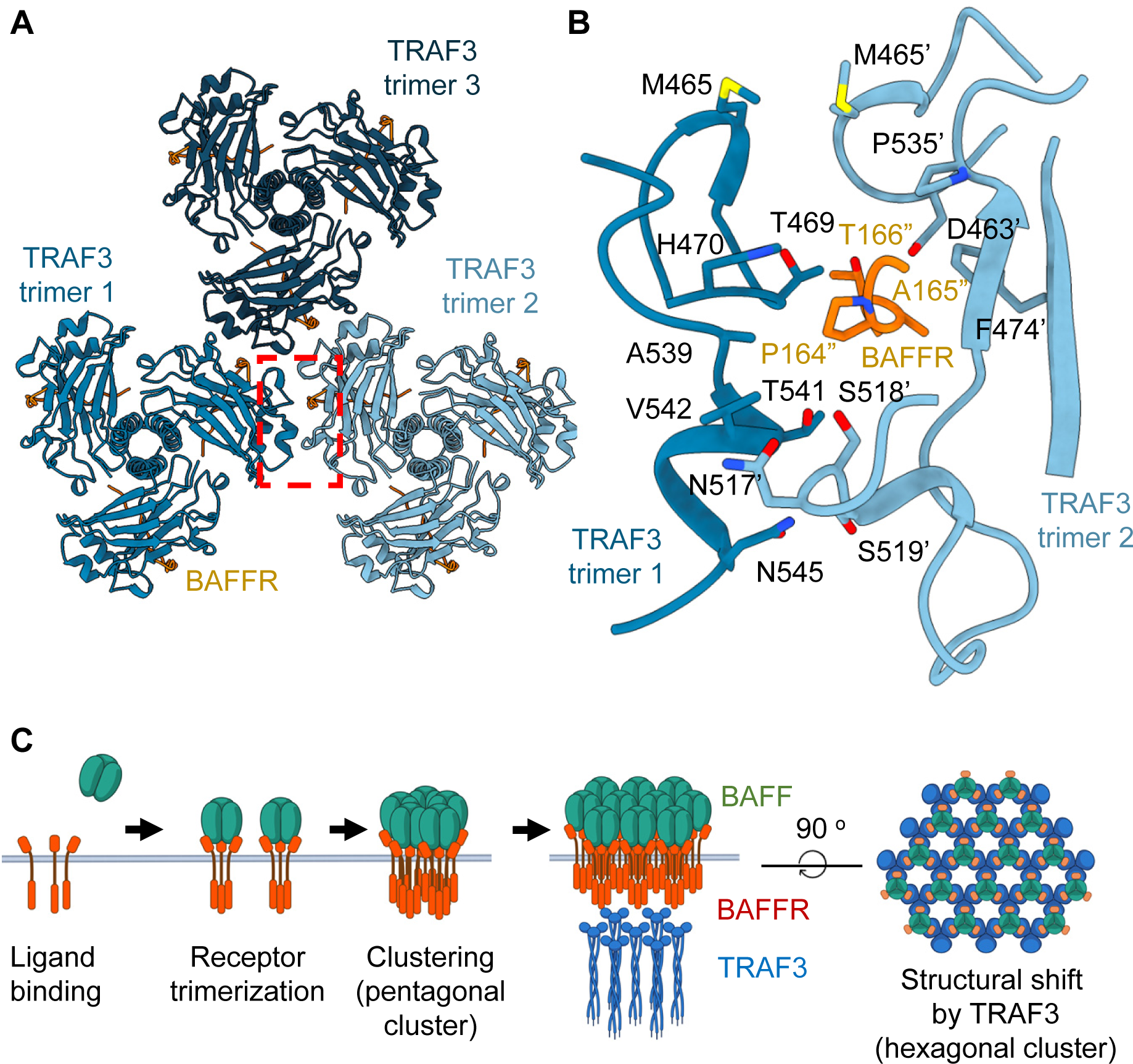
Close-up view of the hisBAFF-BAFFR-TRAF3 cluster and the proposed model of BAFFR activation. (A) Assembly of the three hisBAFF-BAFFRfl-TRAF3 trimers in the cluster. A clustering interface formed by the TRAF3 and BAFFR intracellular domain is highlighted with a red box and zoomed in on (B). (B) A close-up view of a clustering interface. Residue numbers for the second TRAF3 trimer are indicated with an apostrophe. Residue numbers for the BAFFR intracellular domain are shown in orange and indicated with a double apostrophe. (C) The proposed model for the BAFFR activation mechanism.

## DISCUSSION

We used the lipid monolayer method to show that the TNFα-TNFR1 complex formed binary, bent, trigonal, and linear quadruple clusters. We found that dimeric interactions mediate cluster formation between TNFR1 proteins in neighboring TNFα-TNFR1 trimers and that a non-competitive TNFR1 antagonist inhibits cluster formation. We also found that the BAFF-BAFFR complex forms pentagonal, double-pentagonal, and half-sphere clusters when attached to the lipid layer. TRAF3 binds to this cluster and induces a structural shift towards a flat hexagonal cluster. Furthermore, we observed that BAFF mutations that interfere with BAFFR intracellular signaling disrupt receptor clustering. These data demonstrate that highly ordered TNFα-TNFR1 and BAFF-BAFFR clustering plays a crucial role in receptor activation.

The trimerization and expanding network hypotheses are two models proposed to explain how TNFR1 is activated upon binding to the TNFα ligand ^37^. The trimerization hypothesis proposes that the juxtaposition of the three receptors, resulting from the binding of a single TNFα trimer, initiates cell signaling. In this model, the TNFα trimer binds to three receptor molecules, one at each of three TNF monomer-monomer interfaces. The trimerization of the receptors enhances the binding affinity of TRADD and other downstream adaptors to the receptors, ultimately activating signaling pathways ^7^. In contrast, the expanding network hypothesis proposes that the binding of the TNFα trimer to preformed receptor dimers generates a hexagonal array of ligand-receptor complexes. In this model, each TNFα trimer engages three receptor dimers, subsequently triggering network expansion and biological responses. The expanding network hypothesis was proposed based on a crystallographic study of receptor dimers ^38^. Our cryo-EM study with membrane-attached receptors provides the first experimental evidence demonstrating that the proposed network expansion hypothesis is essentially correct, although the network expansion is limited and the resulting cluster size is much smaller because of its twisted nature (Figure 2A).

Our structural observations are consistent with the model proposed based on a single-molecule study using fluorescence microscopy. Recently, Karathanasis *et al.* employed quantitative, single-molecule super-resolution microscopy to directly study TNFR1 assembly in native cellular environments and at physiological cell surface abundance ^39^. The study used quantitative single-molecule localization microscopy combined with TIRF illumination to determine the oligomeric state of TNFR1 and found that, in the absence of TNFα, TNFR1 assembles as a mixture of monomers and dimers. Upon binding with TNFα, TNFR1 clusters into trimers and 9-mers. The study also found that the K32 mutation at the CRD1 interface, known as the PLAD interface, results in monomeric TNFR1 only, which exhibits impaired signal activation. These results underscore the importance of TNFR1 dimerization in receptor activation and the initiation of NF-κB signaling. These fluorescence data are consistent with our cryo-EM findings. We found that the bent cluster with 9-mer receptors is the most abundant cluster when bound to the ligand. We also found that receptor dimerization is incomplete and that a significant proportion of receptors are not involved in receptor dimerization and cluster formation.

The lipid monolayer appeared to play two roles in TNFR1 and BAFFR clustering. First, it restrains receptors in a viscous two-dimensional layer, which enhances the lateral interaction between protein molecules owing to its favorable entropic effect ^48,49^. The protein bound to the 2D lipid layer already has low entropy compared to a freely diffusible protein in a 3D solution space. This reduction in dimensionality favors clustering because the entropy lost by clustering is lower for proteins that are already attached to the lipid layer. Second, the confinement of the protein to the 2D lipid layer limits the shape of the receptor-ligand complexes. For example, previous crystallographic studies demonstrated that the ectodomain of the BAFFR-BAFF complex formed a large globular cage ^18,19^. This structure is evidently impossible when receptors with transmembrane domains are constrained to a flat 2D lipid membrane.

Early crystallographic studies of the BAFFR ectodomain and BAFF complex suggested that BAFF forms a globular cage with 60 monomers, ^18,19^ and this large BAFF cage binds to the BAFFR receptors and activates the intracellular pathway by recruiting TRAF3. A mutational study in the “flap” loop region that mediates BAFF cage formation supported this cage hypothesis ^21^. Other studies have challenged this cage hypothesis because BAFF primarily exists as a trimer in the blood rather than as a cage. Our cryo-EM study demonstrated that both the trimeric and cage forms of BAFF become pentagonal clusters after binding to BAFFR in the membrane and that the binding of TRAF3 to the receptor induces a structural shift to a flat hexagonal cluster. The high-resolution structure of the extracellular part of the BAFF-BAFFR cluster when bound to TRAF3 is unknown. However, the low-resolution map of the extracellular part of the hisBAFF-BAFFRfl-TRAF3 complex in the cluster showed a hexagonal array (Figure 6E). We believe that the pentagonal BAFF-BAFFR cluster, with an inter-trimeric angle of 108°, was flattened, and one additional BAFF-BAFFR trimeric unit was inserted into the cluster, forming a hexagonal lattice with a 120° inter-trimeric angle after TRAF3 binding. We believe that the hexagonal extracellular BAFF-BAFFR cluster structure is more flexible than the pentagonal cluster and that its electron density map fades during map refinement and resolution enhancement using CryoSPARC software ^50^.

The simple aggregation of receptors in close proximity without an ordered structure is insufficient to activate TNFR1 and BAFFR. Although the E223K mutation in BAFF disrupts the ordered structure of the cluster, the mutant ligand-receptor complexes can still aggregate, as shown in Figure 5. Similarly, the binding of the DOM1h nanobody weakens TNFR1 receptor dimerization. However, 40% of TNFR1 receptors on the lipid layer still formed dimers, albeit with a distorted structure (Figure 3B and C). Therefore, the receptor cluster requires a precise structure to activate TNFR1 and BAFFR signaling pathways. A simple, unstructured assembly of receptors does not activate the receptor. However, it remains unclear why a rigid cluster with an ordered structure is required for receptor activation and signal initiation. It is possible that ordered clusters with specific structures have a higher affinity for signaling adaptors, such as TRAF3 or TRADD. Previous crystallographic studies have shown that the N-terminal zinc finger domains of TRAF2 and 6 dimerize ^34,35^. These studies proposed that the dimerization of the TRAF N-terminal domains cross-links the trimeric TRAF C-terminal domains and induces the hexagonal clustering of full-length TRAFs ^36^. Thus, this preformed hexagonal cluster of TRAF3 may have a higher affinity for the BAFF-BAFFR cluster. Alternatively, the clustering of TRAF may be necessary for the proper engagement of downstream effectors, such as NIK, TRAF2, and cIAP, among other signaling proteins. Further research is required to clarify the exact roles of the ordered clustering of TNFR1 and BAFFR receptors.

Nineteen of the 29 TNF receptor family proteins engage with various TRAFs, including TRAF2, 3, and 6, when activated by TNF ligand family proteins ^3^. Most of the remaining TNF receptors use adaptors containing death domains for signaling. Further research is required to determine whether TRAF family proteins other than TRAF3 and death domain-containing adaptors, including TRADD and FADD, form ordered clusters when bound to ligand-receptor clusters. An extensive body of evidence suggests that higher-order clustering is essential for activating single transmembrane receptors other than the TNF receptor family ^51–54^. Furthermore, some GPCRs and ion channels have been proposed to form higher-order complexes ^55–59^. However, most of these clusters are unstable when isolated from the membrane environment; consequently, high-resolution structural characterization of these clusters has been proven difficult. We demonstrated that the lipid monolayer method is a valuable tool for studying the membrane-bound states of these proteins. It provides a large and flat lipid layer that allows protein clustering. Furthermore, it contains only one lipid layer that can bind proteins. This greatly simplifies the cryo-EM analysis of protein clusters. For liposomes, bicelles or nanodiscs containing lipid bilayers, the protein clusters will be formed on both sides of the lipid layers. Deconvoluting images of these two clusters with heterogeneous sizes and shapes will not be trivial.

In conclusion, we demonstrated that the TNF receptor family proteins TNFR1 and BAFFR form highly ordered clusters upon ligand binding. The clustering of receptor-ligand complexes with precise structures is crucial for the activation of intracellular signaling. We propose that other TNF-family receptors have similar mechanisms of action. Structural studies on these receptor clusters will promote the discovery of novel therapeutic agents that can regulate receptor activity by modifying their clustered structures.

## SUPPLEMENTAL INFORMATION

Document S1. Figures S1-S5 Table S1

Table S2 Methods S1

## ACKNOWLEDGMENTS

This work was funded by the National Research Foundation of Korea (RS-2023-00260454 and RS-2024-00344154) and the Technology Innovation Program, MOTIE of Korea (20019707).

## AUTHOR CONTRIBUTIONS

J-O.L. conceived of and supervised the project. C.S.L. expressed and purified the protein, prepared the sample for the cryo-EM study, collected and processed cryo-EM data, and built and refined the atomic models. J.L. conducted all experiments associated with the DOM1h nanobody. C.S.L., J.W.K., and J-O.L. prepared and edited the manuscript with input from all the authors.

## STAR METHODS

## KEY RESOURCES TABLE

| REAGENT or RESOURCE | SOURCE | IDENTIFIER |
| --- | --- | --- |
| Bacterial and virus strains |  |  |
| DH5α | Thermo Fisher Scientific | Cat# EC0112 |
| DH10Bac | Thermo Fisher Scientific | Cat# EC0113 |
| Chemicals, peptides, and recombinant proteins |  |  |
| BestBac™ 2.0 Linearized Baculovirus DNA | Expression Systems | Cat# 91-002 |
| CellFectin II | Gibco™ | Cat# 10362100 |
| Sodium butyrate | Sigma-Aldrich | Cat# 303410-500G |
| Tris-HCl | LPS solution | Cat# TRI05 |
| NaCl | LPS solution | Cat# NACL05 |
| Phenylmethyl fluoride | Sigma-Aldrich | Cat# 52332-25GM |
| Benzamidine chloride | Sigma-Aldrich | Cat# B6506-25G |
| Leupeptin Hemisulfate | GoldBio | Cat# L-010-5 |
| Aprotinin | GoldBio | Cat# A-655-25 |
| Pepstatin | GoldBio | Cat# P-020-25 |
| Imidazole | LPS solution | Cat# IMID500 |
| Thrombin | Lee Biosolutions | Cat# 528-50H |
| PreScission protease | Cytiva | Cat# GE27-0843-01 |
| Fos-Choline-14 (FC-14) | Anatrace | Cat# F312S |
| Glyco-diosgenin (GDN) | Anatrace | Cat# GDN101 |
| 16:0-18:1 PC (POPC) | Avanti Polar Lipids | Cat# 850457 |
| 18:1 DGS-NTA(Ni<sup>2+</sup>) | Avanti Polar Lipids | Cat# 790404 |
| Chloroform | Sigma-Aldrich | Cat# C2432-500ML |
| Acetone | Sigma-Aldrich | Cat# 270725-2L |
| Ethanol | Supelco | Cat# 1.00983 |
| FreeStyle™ 293 Expression Medium | Gibco™ | Cat# 12338018 |
| Sf-900™ II SFM | Gibco™ | Cat# 10902088 |
| ESF 921 Insect Cell Culture Medium | Expression Systems | Cat# 96-001-01 |
| Fetal Bovine Serum | Sigma-Aldrich | Cat# F7524 |
| Deposited data |  |  |
| Coordinates of the TNFα-TNFR1ecto binary cluster | This paper | PDB ID 8ZUI |
| Cryo-EM map of the TNFα-TNFR1ecto binary cluster | This paper | EMD-60484 |
| Cryo-EM map of the TNFα-TNFR1ecto bent cluster | This paper | EMD-60489 |
| Cryo-EM map of the TNFα-TNFR1ecto trigonal cluster | This paper | EMD-60497 |
| Cryo-EM map of the TNFα-TNFR1ecto linear quadruple cluster | This paper | EMD-60491 |
| Cryo-EM map of the TNFα-TNFR1fl binary cluster | This paper | EMD-60490 |
| Cryo-EM map of the TNFα-TNFR1ecto-DOM1h | This paper | EMD-60549 |
| Cryo-EM map of the TNFα-TNFR1ecto-DOM1h on a lipid monolayer | This paper | EMD-60494 |
| Coordinates of the BAFF-BAFFRecto pentagonal cluster | This paper | PDB ID 8ZUJ |
| Cryo-EM map of the BAFF-BAFFRecto pentagonal cluster | This paper | EMD-60485 |
| Cryo-EM map of the BAFF-BAFFRecto double-pentagonal cluster | This paper | EMD-60492 |
| Cryo-EM map of the BAFF-BAFFRecto half-sphere cluster | This paper | EMD-60493 |
| Cryo-EM map of the BAFF-BAFFRfl pentagonal cluster | This paper | EMD-60487 |
| Coordinates of the hisBAFF-BAFFR-TRAF3 cluster | This paper | 8ZUK |
| Cryo-EM map of the hisBAFF-BAFFR-TRAF3 cluster | This paper | EMD-60486 |
| Cryo-EM map of the BAFF-BAFFR-hisTRAF3 cluster | This paper | EMD-60488 |
| Experimental models: Cell lines |  |  |
| Sf9 insect cell | Thermo Fisher Scientific | Cat# 11496015 |
| High Five insect cell | Thermo Fisher Scientific | Cat# B85502 |
| HEK293 GnTI<sup>-</sup> cell | ATCC | Cat# CRL-3022 |
| Recombinant DNA |  |  |
| TNFα in the pAcGP67a vector | This paper | N/A |
| TNFR1ecto in the pAcGP67a vector | This paper | N/A |
| TNFR1fl in the pEG BacMam vector | This paper | N/A |
| DOM1h in the pAcGP67a vector | This paper | N/A |
| BAFF in the pAcGP67a vector | This paper | N/A |
| BAFF(E223K) in the pAcGP67a vector | This paper | N/A |
| BAFF(H218A) in the pAcGP67a vector | This paper | N/A |
| BAFF(K216E) in the pAcGP67a vector | This paper | N/A |
| BAFFRecto in the pAcGP67a vector | This paper | N/A |
| BAFFRfl in the pEG BacMam vector | This paper | N/A |
| TRAF3 in the pVL1393 vector | This paper | N/A |
| hisTRAF3 in the pVL1393 vector | This paper | N/A |
| ALFA nanobody in the pET28a vector | Götzke *et al.* | N/A |
| Software and algorithms |  |  |
| EPU software for Single Particle Analysis | Thermo Fisher Scientific | https://www.thermofisher.com/ |
| CryoSPARC v4.4.1 | Punjani *et al.* | https://cryosparc.com/ |
| Topaz | Bepler *et al.* | https://cb.csail.mit.edu/cb/topaz/ |
| DeepEMhancer | Sanchez-Garcia *et al.* | https://github.com/rsanchezgarc/deepEMhancer |
| Phenix v1.20 | Liebschner *et al.* | https://phenix-online.org/ |
| MolProbity | Williams *et al.* | http://molprobity.biochem.duke.edu/ |
| Coot v0.9.6 | Emsley *et al.* | https://www2.mrc-lmb.cam.ac.uk/personal/pemsley/coot/ |
| UCSF ChimeraX v1.7 | Meng *et al.* | https://www.cgl.ucsf.edu/chimerax/ |
| Other |  |  |
| NHS-activated Sepharose 4 Fast Flow | Cytiva | Cat# 17090601 |
| GST-Sepharose | Cytiva | Cat# 17075601 |
| Superdex 200 increase 10/300 GL gel filtration column | Cytiva | Cat# 28990944 |
| QUANTIFOIL R 1.2/1.3 Gold Grids, size 300 mesh | Jena Bioscience | Cat# X-101-Au300 |
| QUANTIFOIL R 1.2/1.3 Copper Grids, size 300 mesh | Jena Bioscience | Cat# X-101-Cu300 |
| Teflon lipid monolayer preparation chamber | This study | N/A |
| Micro Syringe Pipette | Hamilton | Cat# 84250 |
| AMICON MWCO 50 kDa | Millipore | Cat# UFC505008 |
| AMICON MWCO 10 kDa | Millipore | Cat# UFC901008 |

## RESOURCE AVAILABILITY

### Lead contact

Further information and requests for resources and reagents will be directed and fulfilled by Jie-Oh Lee, the lead contact.

### Materials availability

Plasmids generated in this study are freely available on request.

### Data and code availability

λ The cryo-EM maps and the atomic coordinates were deposited in the Electron Microscopy Data Bank and the Protein Data Bank. Their code numbers are summarized in Supplementary Table S2. This paper does not report the original code.
λ Any additional information required to reanalyze the data reported in this paper is available from the lead contact upon request.

## EXPERIMENTAL MODEL AND SUBJECT DETAILS

DH10Bac (Invitrogen) was cultured in lysogeny broth (LB) at 37℃ to amplify plasmids.

Expi293F cells (Gibco, A14527) were cultured in Expi293 Expression Medium (Thermo Fisher, A1435101) at 37℃ and 5% CO2.

HEK293 GnTI^-^ cells (ATCC, CRL-3022) were cultured in FreeStyle 293 Medium (Gibco, 12338026) Sf9 cells (Gibco, 11496015) were cultured in ESF 921 Medium (Expression System) at 27℃

High Five cells (Thermo Fisher, B85502) were cultured in ESF 921 Medium (Expression System) at 27℃

## METHOD DETAILS

### Expression and purification of soluble TNF*α*

The gene encoding the ectodomain of human TNFα, amino acids 77V∼233L, with a thrombin cleavage site and a hexa-histidine tag at the C-terminus was synthesized (Twist Bioscience) and cloned into a pAcGP67a baculovirus transfer vector (BD Biosciences) (Table S1). Recombinant baculovirus was generated by co-transfection with a linearized baculovirus genome, BestBac2.0 (Expression Systems), into Sf9 insect cells. Protein was expressed in High Five insect cells cultured in ESF 921 media (Expression Systems) by adding 4% (v/v) baculovirus at 27℃ for 72 hours. The secreted protein was bound to cOmplete His-Tag Purification Resin (Roche) and eluted using an elution buffer containing 20 mM Tris-HCl, 150 mM NaCl, and 300 mM imidazole at pH 7.8. The protein was further purified by a Superdex 200 increase 10/300 GL gel filtration column (Cytiva) equilibrated with a buffer containing 20 mM Tris-HCl and 150 mM NaCl at pH 7.8. For TNFα without the histidine tag, protease cleavage and additional purification steps were taken. Thrombin treatment was carried out at 4°C for 16 hours, followed by another round of gel filtration chromatography. The solution is then passed through the His-Tag Purification column to remove un-cleaved protein.

### Expression and purification of TNFR1

The gene encoding the ectodomain of human TNFR1, amino acid numbers 1L∼182T, with an octa-histidine tag at the C-terminus, was synthesized (Twist Bioscience) and cloned into a pAcGP67a baculovirus transfer vector (BD Biosciences) (Table S1). Recombinant baculovirus was generated by co-transfection with a linearized baculovirus genome, BestBac2.0 (Expression Systems), into Sf9 insect cells. Protein was expressed in High Five insect cells cultured in ESF 921 media (Expression System) by adding 4% (v/v) baculovirus at 27°C for 72 hours. The secreted protein was bound to cOmplete His-Tag Purification Resin (Roche) and eluted using an elution buffer containing 20 mM Tris-HCl, 150 mM NaCl, and 300 mM imidazole at pH 7.8. The protein was further purified by a Superdex 200 increase 10/300 GL gel filtration column (Cytiva) equilibrated with a buffer containing 20 mM Tris-HCl and 150 mM NaCl at pH 7.8.

The gene encoding the full-length human TNFR1 was synthesized (Twist Bioscience) and cloned into a pEG BacMam baculovirus transfer vector (Table S1) ^60^. PreScission protease site, mCherry, thrombin cleavage site, and ALFA tag were added to the C-terminus of TNFR1. The plasmid was transformed into DH10Bac *E.coli* for transposition into the bacmid ^60^. The recombinant baculovirus was generated by transfecting Sf9 insect cells with the bacmid DNA. Protein was expressed in HEK293 GnTI^-^ cells cultured in the FreeStyle 293 media (Thermo Fisher Scientific) supplemented with 1% (v/v) fetal bovine serum (Sigma-Aldrich) in a 10% CO_2_ shaking incubator. The cells were infected by treating 8% (v/v) baculovirus at a density of 3×10^6^ cells per ml. Protein expression was enhanced by treating with 10 mM sodium butyrate (Sigma-Aldrich) after 12-16 hours of infection and further incubated for 48 to 60 hours. The cells were collected by centrifugation at 7,261g for 30 min, flash-frozen in liquid nitrogen, and stored at −80°C.

For purification of the full-length TNFR1, a frozen aliquot of the cells was thawed and resuspended in a buffer, 20 mM Tris-HCl pH 8.0, 150 mM NaCl, 0.5 mM phenylmethyl fluoride (PMSF), 1 mM benzamidine chloride, 1 μg/ml leupeptin, 1 μg/ml aprotinin, and 1 μg/ml pepstatin and lysed using a homogenizer. The membrane fraction was collected for one hour by ultracentrifugation, 200,000g. The pellet was resuspended in the lysis buffer, and the membrane fraction was solubilized by adding 1% (w/v) Fos-Choline-14 (FC-14, Anatrace) and incubated for one hour at 4°C with gentle rotation. The insoluble fraction was removed by ultracentrifugation at 200,000g for one hour. The supernatant was loaded onto a column packed with an agarose resin conjugated to an anti-ALFA nanobody ^61^. The anti-ALFA nanobody was conjugated to NHS-activated Sepharose 4 resin (Cytiva) following the manufacturer’s protocol. A total of 20 µmol of ALFA nanobody was conjugated to 1 ml of NHS-activated Sepharose 4 resin, and unreacted NHS resin was blocked using 20 mM Tris-HCl pH 8.0. The anti-ALFA-resin was washed with 40 column volumes of wash buffer containing 200 mM Tris-HCl pH 8.0, 150 mM NaCl, and 0.005% FC-14. Protein was eluted by treating with 1% PreScission Protease (Cytiva) overnight. The eluted protein solution was concentrated using an ultracentrifugal filter with a 50 kDa cutoff (Merck Millipore).

### Expression and purification of the wild type and the mutant of BAFF

The gene encoding the ectodomain of human BAFF, amino acids 134A∼285L, with a six-histidine tag and a thrombin cleavage site at the N-terminus, was synthesized (Twist Bioscience) and cloned into a pAcGP67a baculovirus transfer vector (BD Biosciences) (Table S1). Recombinant baculovirus was generated by co-transfection with a linearized baculovirus genome, BestBac2.0 (Expression Systems), into Sf9 insect cells. Protein was expressed in High Five insect cells cultured in ESF 921 media (Expression Systems) by adding 4% (v/v) baculovirus at 27℃ for 72 hours. The secreted protein was bound to cOmplete His-Tag Purification Resin (Roche) and eluted using an elution buffer containing 20 mM Tris-HCl, 150 mM NaCl, and 300 mM imidazole at pH 7.8. The histidine tag was removed by thrombin incubation at 4℃ for 16 hours. For purification of the trimeric BAFF, the cleaved protein was purified, without concentration, by a Superdex 200 increase 10/300 GL gel filtration column (Cytiva) equilibrated with a buffer containing 20 mM Tris-HCl and 150 mM NaCl at pH 7.8. The fractions with molecular weight corresponding to the trimeric BAFF were collected (Figure S3A). For purification of the cage form of the BAFF, the cleaved protein was concentrated to higher than 1.5 mg/ml using ultrafiltration with a membrane with molecular weight cutoff of 50 kDa and purified by a Superdex 200 increase 10/300 GL gel filtration column (Cytiva) equilibrated with a buffer containing 20 mM Tris-HCl and 150 mM NaCl at pH 7.8. The fractions corresponding to the BAFF cage were collected.

### Expression and purification of BAFFR

The gene encoding the ectodomain of human BAFFR, amino acids 2R∼78L, with a FLAG tag at the N-terminus and an octa-histidine tag at the C-terminus, was synthesized (Twist Bioscience) and cloned into a pAcGP67a baculovirus transfer vector (BD Biosciences) (Table S1). Recombinant baculovirus was generated by co-transfection with a linearized baculovirus genome, BestBac2.0 (Expression Systems), into Sf9 insect cells. Protein was expressed in High Five insect cells cultured in ESF 921 media (Expression Systems) by adding 4% (v/v) baculovirus at 27℃ for 72 hours. The secreted protein was bound to cOmplete His-Tag Purification Resin (Roche) and eluted using an elution buffer containing 20 mM Tris-HCl, 150 mM NaCl, and 300 mM imidazole at pH 7.8. The protein was further purified by a Superdex 200 increase 10/300 GL gel filtration column (Cytiva) equilibrated with a buffer containing 20 mM Tris-HCl and 150 mM NaCl at pH 7.8.

The gene encoding the full-length human BAFFR was synthesized (Twist Bioscience) and cloned into a pEG BacMam baculovirus transfer vector (Table S1). A PreScission protease site, mCherry, thrombin cleavage site, and ALFA tag were added to the C-terminus of TNFR1. The plasmid was transformed into DH10Bac *E.coli* for transposition into the bacmid ^60^. The recombinant baculovirus was generated by transfecting Sf9 insect cells with the bacmid DNA. Protein was expressed in HEK293 GnTI^-^ cells cultured in the FreeStyle 293 media (Thermo Fisher Scientific) supplemented with 1% (v/v) fetal bovine serum (Sigma-Adrich) in a 10% CO_2_ shaking incubator. The cells were infected by treating 8% (v/v) baculovirus at a density of 3×10^6^ cells per ml. Protein expression was enhanced by treating with 10 mM sodium butyrate (Sigma-Aldrich) after 12-16 hours of infection and further incubated for 48 to 60 hours. The cells were collected by centrifugation at 7,261g for 30 min, flash-frozen in liquid nitrogen, and stored at −80℃.

For purification of the full-length BAFFR, a frozen aliquot of the cells was thawed and resuspended in a buffer, 20 mM Tris-HCl pH 8.0, 150 mM NaCl, 0.5 mM PMSF, 1 mM benzamidine chloride, 1 μg/ml leupeptin, 1 μg/ml aprotinin, and 1 μg/ml pepstatin and lysed using a homogenizer. The membrane fraction was collected for one hour by ultracentrifugation, 200,000g. The pellet was resuspended in the lysis buffer, and the membrane fraction was bound to the ligand by adding 16 µg/ml of BAFF for one hour and a half at 4℃ with gentle rotation. The BAFF-BAFFRfl complex was extracted with 1% (w/v) FC-14 (Anatrace) and incubated for two hours at 4℃ with gentle rotation. The insoluble fraction was removed by ultracentrifugation at 200,000g for one hour. The supernatant was loaded onto a column packed with an agarose resin conjugated to an anti-ALFA nanobody ^61^. The resin was washed with 40 column volumes of wash buffer containing 20 mM Tris-HCl pH 8.0, 150 mM NaCl, and 0.005% FC-14. Protein was eluted by treating with 1% PreScission Protease (Cytiva) overnight.

### Expression and purification of the truncated TRAF3

The gene encoding the coiled-coil and TRAF-C domains of human TRAF3 with a thrombin cleavage site, a hexa-histidine, and the ALFA tag at the C-terminus was synthesized (Twist Bioscience) and cloned into a pVL1393 baculovirus transfer vector (BD Biosciences) (Table S1). Recombinant baculovirus was generated by co-transfection with a linearized baculovirus genome, BestBac2.0 (Expression Systems), into Sf9 insect cells. Protein was expressed in High Five insect cells cultured in ESF 921 media (Expression Systems) by adding 4% (v/v) baculovirus at 21℃ for 72 hours. The cells were collected by centrifugation at 7,261g for 30 minutes, flash-frozen in liquid nitrogen, and stored at −80℃.

For purification, a frozen aliquot of the cells was thawed and resuspended in a buffer, 20 mM Tris-HCl pH 8.0, 150 mM NaCl, 0.5 mM PMSF, and lysed using a microfluidizer (Microfluidics). The protein was bound to cOmplete His-Tag Purification Resin (Roche) and eluted using an elution buffer containing 20 mM Tris-HCl, 150 mM NaCl, and 300 mM imidazole at pH 7.8. The protein was further purified by a Superdex 200 increase 10/300 GL gel filtration column (Cytiva) equilibrated with a buffer containing 20 mM Tris-HCl and 150 mM NaCl at pH 7.8. For protein purification without the histidine tag, the protein eluted from the cOmplete His-Tag Purification Resin was treated with thrombin at 4°C for 16 hours, followed by Supdex-200 gel filtration chromatography. The protein-containing fractions were collected and passed through the His-Tag Purification column to remove uncleaved protein.

### Expression and purification of the DOM1h nanobody

The gene encoding the DOM1h-574-208 nanobody with a thrombin cleavage site and a hexa-histidine tag at the C-terminus was synthesized (Twist Bioscience) and cloned into pAcGP67A baculovirus transfer vector (BD Biosciences) (Table S1). Recombinant baculovirus was generated by co-transfection with a linearized baculovirus genome, BestBac2.0 (Expression Systems), into Sf9 insect cells. Protein was expressed in High Five insect cells cultured in ESF 921 media (Expression Systems) by adding 4% (v/v) baculovirus at 27℃ for 72 hours. The secreted protein was bound to cOmplete His-Tag Purification Resin (Roche) and eluted using an elution buffer containing 20 mM Tris-HCl, 150 mM NaCl, and 300 mM imidazole at pH 7.8. The eluted protein was treated with thrombin at 4°C for 16 hours, followed by a Superdex 200 increase 10/300 GL gel filtration column (Cytiva) equilibrated with a buffer containing 20 mM Tris-HCl and 150 mM NaCl at pH 7.8. The protein-containing fractions were collected and passed through the His-Tag Purification column to remove uncleaved protein.

### Preparation of the lipid monolayer and cryo-EM grids for the TNF*α*-TNFR1 and BAFF-BAFFR complexes

The lipid monolayer was prepared as previously reported ^40^. Briefly, a Teflon block with 85 µl well and a side injection tunnel was prepared (Figure S1A). The wells were washed with ethanol and distilled water. The Teflon block was placed in a humid environment in a Petri dish with wet filter paper. Then, each well was filled with 65 µl of buffer containing 20 mM Tris-HCl and 150 mM NaCl at pH 7.8. One microliter of a lipid solution containing 0.8 mg/ml POPC (Sigma) and 0.2 mg/ml DGS-NTA(Ni^2+^) (Sigma) dissolved in chloroform was gently added on top of the buffer. The Teflon block was left to sit in the Petri dish at room temperature for 30 minutes, allowing chloroform to evaporate. The lipid monolayer forms spontaneously.

For cryo-EM grid preparation, 20 µl of a solution containing the receptor-ligand complex was injected into the side tunnel of the well. Then, the protein solution in the well was gently mixed by a pipet inserted into the side tunnel and incubated at room temperature for 2 hours to allow the histidine-tagged protein to bind to the lipid monolayer. Then, a Quantifoil 1.2/1.3 gold EM grid with the carbon side facing down was placed on top of the well and was left to sit for 30 minutes to allow the lipid layer to adhere to the carbon film. Next, 15 µl of buffer was added to the side tunnel of the well. The grid was gently picked up with forceps and then plunge-frozen using a Vitrobot Mark IV (Thermo Fisher Scientific) under 18°C, 100% humidity, and no force condition after 6 seconds of blotting.

### Preparation of lipid monolayer and cryo-EM grids for the hisBAFF-BAFFRfl-TRAF3 complex

The lipid monolayer was prepared as written in the previous paragraphs. For cryo-EM grid preparation, 20 µl of buffer was removed from the side tunnel of the Teflon well, and 20 µl of 0.1 mg/ml of the full-length BAFFR in complex with BAFF tagged with hexa-histidine was added. The solution was gently mixed with a pipet and incubated at room temperature for two hours. 30 µl of the protein solution in the well was replaced twice by 30 µl of buffer solution to reduce the concentration of the protein unbound to the monolayer. Then, 30 µl of 0.2 mg/ml TRAF3 without the hexa-histidine tag was added to the side tunnel and incubated for five hours at room temperature. A Quantifoil 1.2/1.3 gold EM grid with the carbon side facing down was placed on top of the Teflon well and was left to sit for 30 minutes to allow the lipid layer to adhere to the carbon film. Next, 15 µl of buffer was added to the side tunnel of the well. The grid was gently picked up with forceps and then plunge-frozen using a Vitrobot Mark IV (Thermo Fisher Scientific) under 18°C, 100% humidity, and no force condition after 6 seconds of blotting.

### Preparation of lipid monolayer and cryo-EM grids for the BAFF-BAFFRfl-hisTRAF3 complex

The lipid monolayer was prepared as written in the previous paragraphs. For cryo-EM grid preparation, 20 µl of buffer was removed from the Teflon well, and 20 µl of 0.1 mg/ml of TRAF3 with an N-terminal octa-histidine tag was added. The solution was gently mixed with a pipet and incubated at room temperature for two hours. 30 µl of the protein solution in the well was replaced twice by 30 µl of buffer solution to reduce the protein concentration unbound to the monolayer. Then, 30 µl of 0.2 mg/ml of the full-length BAFFR in complex with BAFF without the hexa-histidine tag was added to the side tunnel of the well and incubated for five hours at room temperature. A Quantifoil 1.2/1.3 gold EM grid with the carbon side facing down was placed on top of the Teflon well and was left to sit for 30 minutes to allow the lipid layer to adhere to the carbon film. Next, 15 µl of buffer was added to the side tunnel of the well. The grid was gently picked up with forceps and then plunge-frozen using a Vitrobot Mark IV (Thermo Fisher Scientific) under 18°C, 100% humidity, and no force condition after 6 seconds of blotting.

### Preparation of cryo-EM grids for the TNF*α*-TNFR1-DOM1h complex

TNFα, TNFR1ecto, and DOM1h proteins were mixed with a 1:1.3:1.5 molar ratio to prepare the protein complex. The total protein concentration was 1.1 mg/ml. 0.00021% glycol-diosgenin (GDN) detergent was added to the buffer solution to reduce the orientation preference of the protein particles in the cryo-EM grids. The 300-mesh Quanti-foil R 1.2/1.3 copper EM grids (Quantifoil Micro Tools) were discharged at 15 mA for 60 seconds using a glow-discharger (PELCO). Then, cryo-EM grid samples were prepared using a Vitrobot Mark IV (Thermo Fisher Scientific) under 4℃ and 100% humidity. Three and a half microliters of the sample were loaded onto the cryo-EM grid and blotted 5 seconds before plunge-freezing. Preparation of cryo-EM grids for the TNFα-TNFR1-DOM1h complex bound to the lipid monolayer was prepared as described in the “Preparation of the lipid monolayer and cryo-EM grids for the TNFα-TNFR1 and BAFF-BAFFR complexes” section.

### Cryo-EM data collection

The data collection statistics are summarized in Table S2. The TNFα-TNFR1ecto, the BAFF-BAFFRecto, and the TNFα-TNFR1ecto-DOM1h data sets were collected using a Titan Krios microscope (Thermo Fisher Scientific) at 300 kV, equipped with an energy filter and a K3 direct electron detector (Gatan) operating in counting mode at 100,500x magnification with a pixel size of 0.82 Å. Each cryo-EM movie was recorded over 49 frames with a total dose of 50 e/Å². The BAFF-BAFFRfl and the hisBAFF-BAFFRfl-TRAF3 data sets were collected using a Titan Krios microscope at 300 kV, equipped with an energy filter and a Falcon4i direct electron detector (Thermo Fisher Scientific) operating in counting mode at 130,000x magnification with a pixel size of 0.94 Å. Each cryo-EM movie was recorded in the Electron Event Representation (EER) format with a total dose of 40 e/Å². The trigonal cluster of TNFα-TNFR1ecto, the TNFα-TNFR1fl, the TNFα-TNFR1ecto-DOM1h (lipid monolayer), the half-sphere cluster of BAFF-BAFFRecto, the BAFF-BAFFRecto mutants, and the BAFF-BAFFRfl-hisTRAF3 data sets was collected using a Talos Glacios microscope at 200 kV, equipped with a Falcon4 detector operating in counting mode at 92,000x magnification with a pixel size of 1.088 Å. All data sets were automatically acquired using EPU software for single particle analysis (Thermo Fisher Scientific).

### Cryo-EM data preprocessing

All cryo-EM data processing was performed using the cryoSPARC v4.4.1 program ^50^. The cryo-EM movie files were preprocessed using the patch motion correction and the patch contrast transfer function (CTF) methods. The EER format data were imported into the CryoSPARC program with an up-sampling factor of 1, followed by patch motion correction and patch CTF estimation. Poor-quality micrographs were manually curated based on the CTF estimations, ice thickness, and the total motion of the frames. Initial particles were picked using the blob-picking method and purified by the 2D class average method. The high-quality 2D class averages were selected for Topaz picking model generation ^62^. The particle picking method is further optimized by repeating 2-3 rounds of Topaz analysis and multi-rounds of 2D classification.

### Data processing for the TNF*α*-TNFR1 complexes

Data processing strategies are summarized in Methods S1. For the binary cluster of the TNFα-TNFR1ecto complex, particles were extracted with a box size of 360 pixels and binned by 120 pixels. The initial model was generated using *Ab-initio* reconstruction with C1 symmetry. The electron density map was refined through homogeneous and non-uniform refinements by imposing C2 symmetry. The final map was obtained by un-binning the particles and performing non-uniform refinement with C2 symmetry. The final map was sharpened, and the noise was reduced using DeepEMhancer, an automatic deep learning-based sharpening method ^63^. A similar processing scheme was used for the binary cluster of the TNFα-TNFR1fl complex data set.

Data processing for the bent cluster of the TNFα-TNFR1ecto complex, particles were extracted with a box size of 512 pixels and binned by 128 pixels. The particles obtained from the 2D class averages included both the bent and binary clusters. To differentiate the two clusters, we further sorted the particles by performing heterogeneous refinement of the bent and binary clusters obtained from multi-class *Ab-initio* reconstruction. Subsequently, the bent cluster particles were selected, and homogeneous and non-uniform refinements were conducted to obtain a refined electron density map.

Data processing for the linear quadruple cluster of the TNFα-TNFR1ecto complex, particles were extracted with a box size of 512 pixels and binned by 128 pixels. The particles were further sorted using multi-class *Ab-initio* reconstruction. After sorting, classes corresponding to the linear quadruple cluster were selected, and the final map was obtained by homogenous refinement by imposing C2 symmetry. Data processing for the trigonal cluster of the TNFα-TNFR1ecto complex, particles were extracted with a box size of 384 pixels. The initial model was generated using *Ab-initio* reconstruction with C3 symmetry and refined through homogeneous and non-uniform refinements with C3 symmetry. The final map was sharpened, and the noise was reduced using DeepEMhancer ^63^.

### Data processing for the TNF*α*-TNFR1ecto-DOM1h complex

Data processing for the TNFα-TNFRecto-DOM1h complex, particles were initially extracted with a box size of 256 pixels and binned by 128 pixels. The initial model was generated using the *Ab-initio* reconstruction method with C1 symmetry, followed by particle re-centering and un-binning. The electron density map was refined through homogeneous and non-uniform refinements with C3 symmetry. Due to the high flexibility and independent motions of the domains, the CRD4 domain of TNFR1ecto and the DOM1h nanobody could not be resolved in the electron density map. To address this issue, we conducted a focused refinement to improve the resolution of the map for the CRD4 and the nanobody regions. The signals from the two CRD4 domains and the nanobodies bound to them were subtracted using the particle subtraction method. The map for the remaining CRD4 domain and the nanobody was enhanced using local refinement with a mask covering the trimeric TNFα ligands and a single TNFR1ecto bound to the nanobody. After model building, a composite map reproducing C3 symmetry covering all three nanobodies was generated using the “Combine Focus Maps” function of the program Phenix ^64^.

Data processing for the binary cluster of the TNFα-TNFRecto-DOM1h complex bound to a lipid monolayer, particles were initially extracted with a box size of 300 pixels and binned by 100 pixels. The initial model was generated using *Ab-initio* reconstruction with C2 symmetry and refined through homogeneous and non-uniform refinements. The structure of the trimeric TNFα-TNFR1ecto complex was fitted into the obtained map, and the initial volume was generated using the molmap function in the ChimeraX program with a 30 Å low-resolution filter ^65^. The final map was obtained using non-uniform refinement with C2 symmetry.

### Data processing for the BAFF-BAFFR clusters

For the pentagonal cluster of the BAFF-BAFFRecto complex, particles were extracted with a box size of 400 pixels. The initial model was generated using the *Ab-initio* reconstruction with C1 symmetry. The electron density map was refined through homogeneous and non-uniform refinements by imposing C5 symmetry. The final map was improved by per-particle-based motion correction using the local motion correction method. A similar scheme was used to process the pentagonal cluster of the BAFF-BAFFRfl data set.

For the double pentagonal cluster of the BAFF-BAFFRecto complex, initial particles were extracted with a box size of 500 pixels. The initial model generation using the *Ab-initio* reconstruction method failed, presumably due to the limited view of particles or the flexible nature of the complex. Therefore, we generated the initial model using the double pentagonal substructure extracted from the crystal structure of the full-size BAFF-BAFFR cage with PDB ID 4V46 ^18^. The initial volume was generated using the molmap function in the ChimeraX program with a 30 Å low-resolution filter ^65^. By performing homogeneous refinement with C2 symmetry, the final map was obtained with a nominal resolution of 3.5 Å.

Data processing for the half-spherical cluster of the BAFF-BAFFRecto complex, initial particles were extracted with a box size of 400 pixels and binned by 100 pixels. Several 2D class averages showed convolutions with other oligomeric states, and therefore, particles were sorted by performing heterogeneous refinement. The initial models for heterogeneous refinement were generated by the *Ab-initio* reconstruction using a selected subset of particles. The electron density map was refined through homogeneous and non-uniform refinements by imposing C3 symmetry. The final map was improved by per-particle-based motion correction using the local motion correction method.

### Data processing for the BAFF-BAFFRfl-TRAF3 clusters

Data processing for the hisBAFF-BAFFRfl-TRAF3 cluster, initial particles were extracted with a box size of 700 pixels and binned by 200 pixels. Duplicated particles were removed during particle picking and 2D classification by setting a minimum separation distance of 20 Å. The initial model was generated using the *Ab-initio* reconstruction with C1 symmetry. The electron density map was refined through homogeneous refinement by imposing C3 symmetry. The map quality of the region near TRAF3 was enhanced by conducting homogeneous refinement with a mask covering the seven TRAF3 trimers. The final map was sharpened, and the noise was reduced using DeepEMhancer ^63^.

Data processing for the BAFF-BAFFRfl-hisTRAF3 cluster, initial particles were extracted with a box size of 400 pixels and binned by 200 pixels. Particle sorting in three dimensions and generation of the initial model was performed using the *Ab-initio* reconstruction with C1 symmetry. Low-quality particles were removed, and the initial model, containing TRAF3 in the center, was used for further refinement. The refined electron density map was obtained by conducting homogeneous and non-uniform refinements with C3 symmetry. Seven TRAF3 units were fitted into the obtained map, and the initial volume was generated using the molmap function in the ChimeraX program with a 10 Å low-resolution filter. The final map was obtained using non-uniform refinement with a mask covering the seven TRAF3 trimers in the center.

### Model building

The initial structural templates for the binary cluster of the TNFα-TNFR1ecto complex were obtained from crystallographic structures of TNFα with PDB ID 7KPB ^66^ and TNFR1ecto with PDB ID 7KP7 ^8^. These initial models were fitted into the cryo-EM density map using ChimeraX ^65^. The resulting structure of the binary cluster of the TNFα-TNFR1ecto complex was refined through multiple rounds of model building using Coot and Phenix ^64,67^. The bent, trigonal and linear quadruple clusters of the TNFα-TNFR1ecto and the binary cluster of the TNFα-TNFRfl complexes were modeled by the rigid-body fitting of the trimeric TNFα-TNFR1ecto structure to the map. The TNFα-TNFR1ecto-DOM1h complex was modeled with a similar method. The initial model for the DOM1h nanobody was generated using AlphaFold 3 ^68^.

The initial model for the pentagonal cluster of the BAFF-BAFFRecto complex was obtained by rigid-body fitting of the pentagonal substructure extracted from the crystal structure of the BAFF-BAFFRecto cage with PDB ID 4V46. The resulting model was refined through multiple rounds of model building using Coot and Phenix with reference model restraints. The structures of the double pentagonal and half-spherical clusters of the BAFF-BAFFRecto complex and the pentagonal cluster of the BAFF-BAFFRfl complex were initially built by the rigid-body fitting of the trimeric BAFF-BAFFRecto complex and refined by a similar method.

The initial model for the hisBAFF-BAFFRfl-TRAF3 complex was generated by combining previously reported structures of TRAF3, PDB ID 8T5P ^69^, and the BAFFR intracellular domain, PDB ID 2GKW ^26^. The model was refined through iterative model-building steps using the Coot and Phenix programs. The reported crystal structure of TRAF3 contains a shorter coiled-coil domain than the TRAF3 protein we used for the cryo-EM study. The missing part of the coiled-coil alpha helix was extracted from a TRAF3 model predicted by AlphaFold 3 ^68^. The refinement statistics are summarized in Table S2. All figures with electron density maps, structural analyses, and representations were generated using ChimeraX and Coot ^65,67^.

## Supplemental information

**Figure S1.**
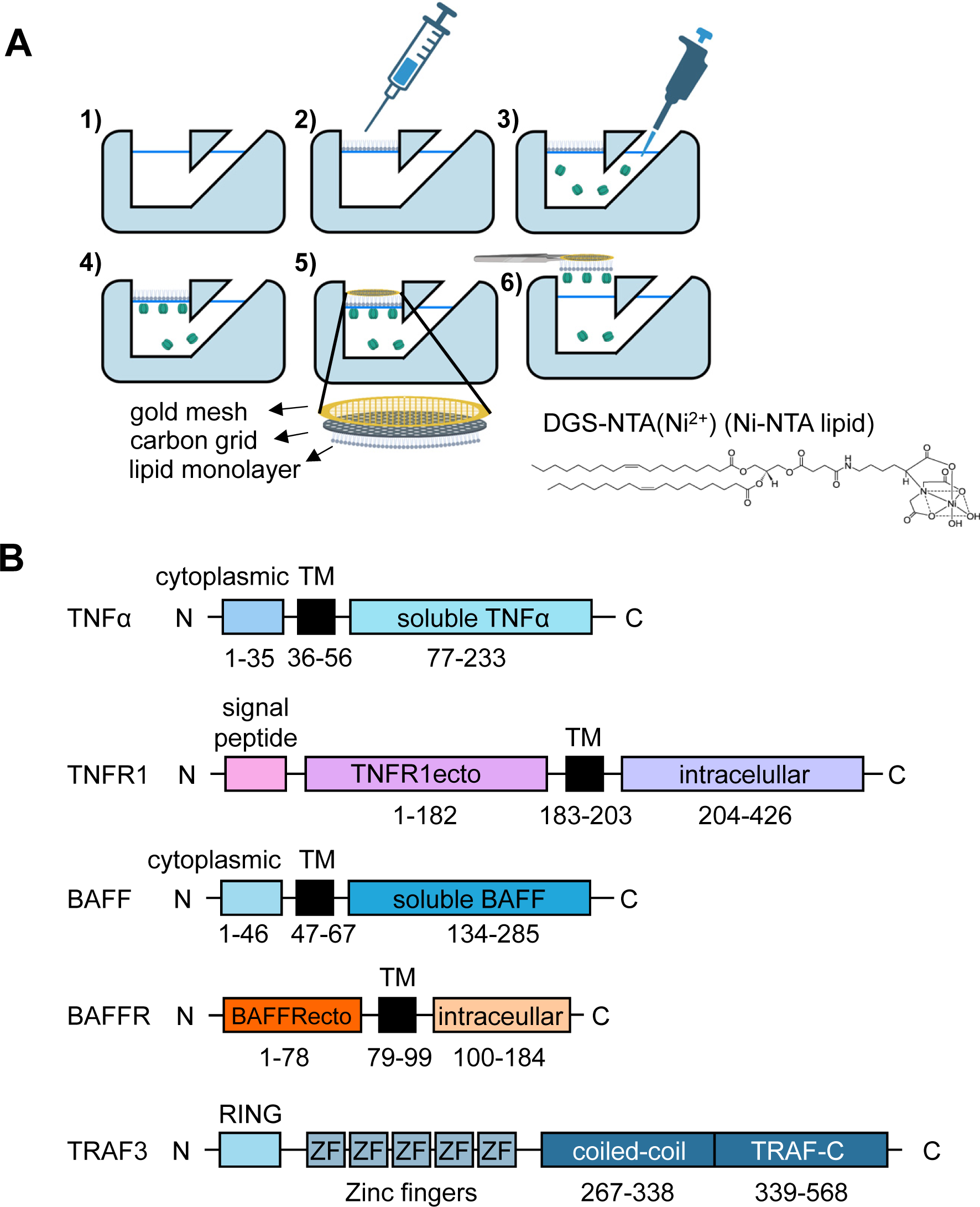
Preparation of the lipid monolayer grids and summary of the protein sequences, related to Figure 1. (A) Grid preparation procedure: 1) addition of the buffer to the chamber; 2) addition of the lipid-chloroform drop to the buffer surface followed by chloroform evaporation; 3) sample injection; 4) attachment of the protein to the Ni-NTA lipid monolayer; 5) attachment of the lipid layer to the grid via hydrophobic interactions; and 6) grid transfer for cryo-EM analysis. The chemical structure of the DGS-NTA(Ni^2+^) is shown. (B) Schematic diagrams showing the protein domain organizations.

**Figure S2.**
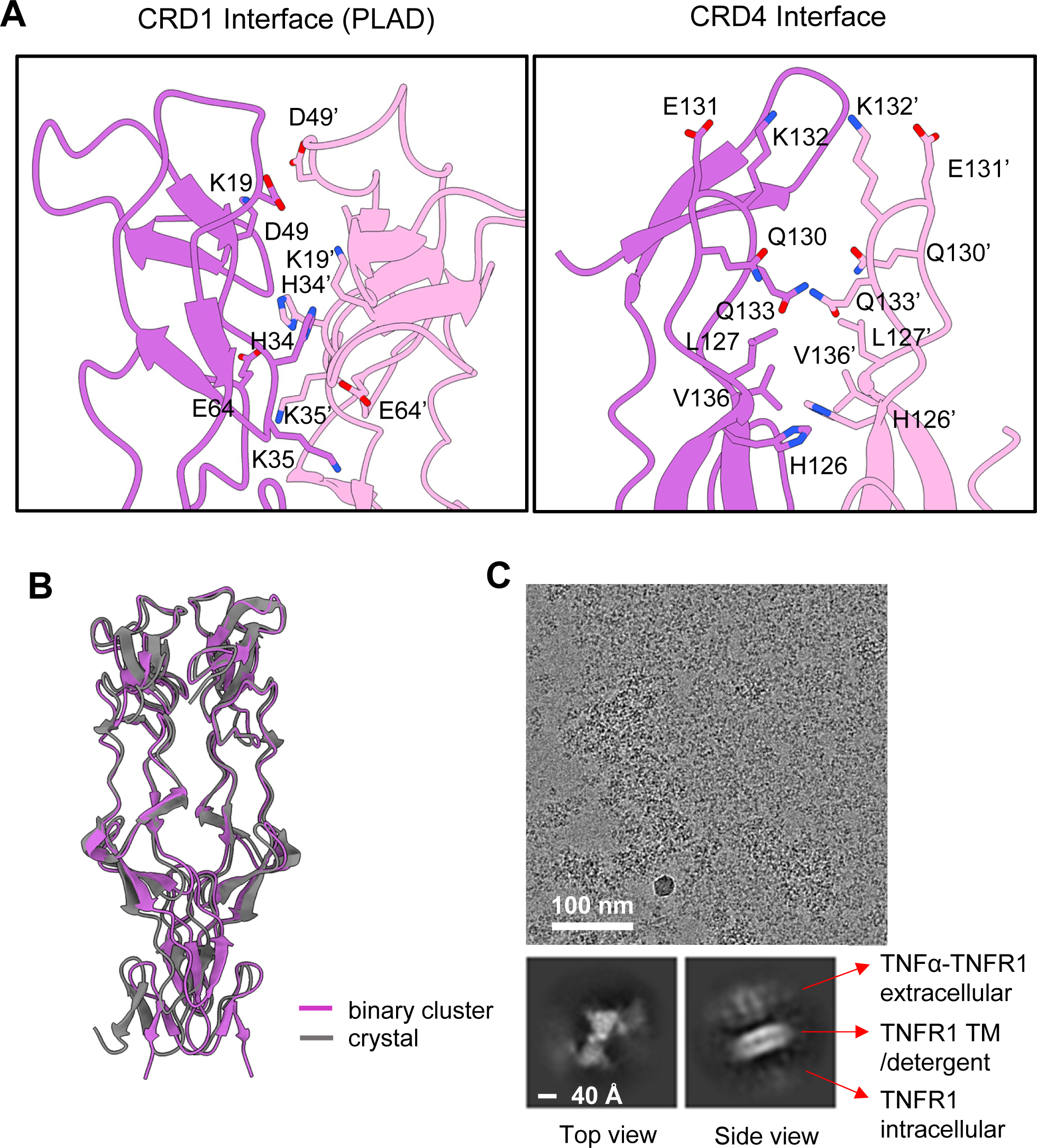
Structure of the TNFR1 dimer in the cluster, related to Figure 2. (A) Close-up view of the CRD1 and CRD4 interfaces between the dimerized TNFR1 receptors in the binary cluster. The view corresponds to that of the lower panel in Figure 2A. The residue numbers of the second TNFR1 are indicated with apostrophes. (B) Structural comparison of the TNFR1 dimer in the binary cluster and the previously reported crystal structure (PDB code: 1NCF). (C) A representative cryo-EM image (upper) and the 2D class averages of the TNFα-TNFR1fl complex (lower two panels).

**Figure S3.**
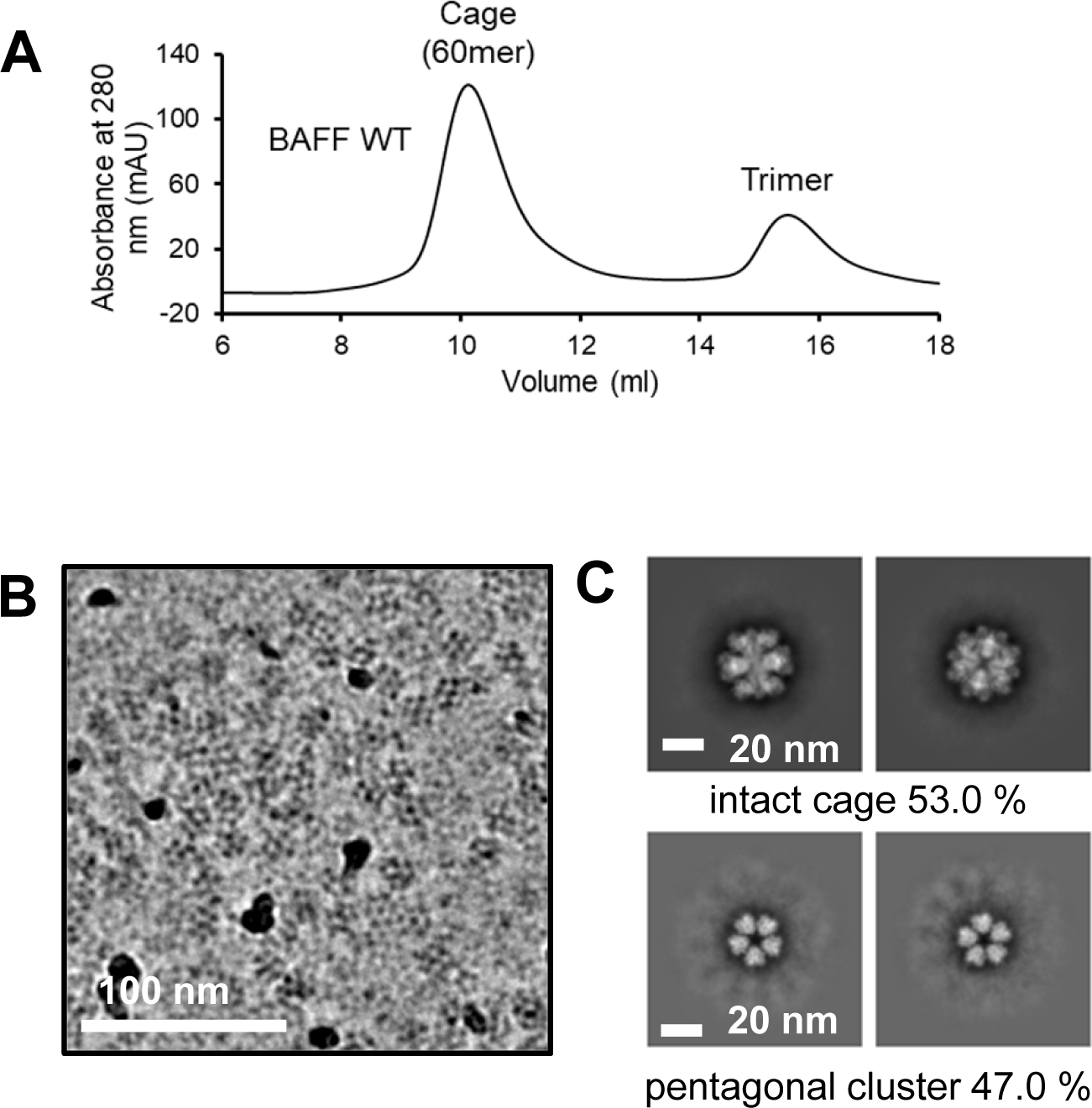
Dissociation of the BAFF cage into the pentagonal cluster by binding to BAFFRecto attached to the lipid layer, related to Figure 4. (A) Preparation of the trimeric and cage forms of BAFF. The protein samples were analyzed using Superdex-200 gel filtration chromatography. (B) A representative cryo-EM image. (C) 2D class averages of the intact cages and dissociated pentagonal clusters.

**Figure S4.**
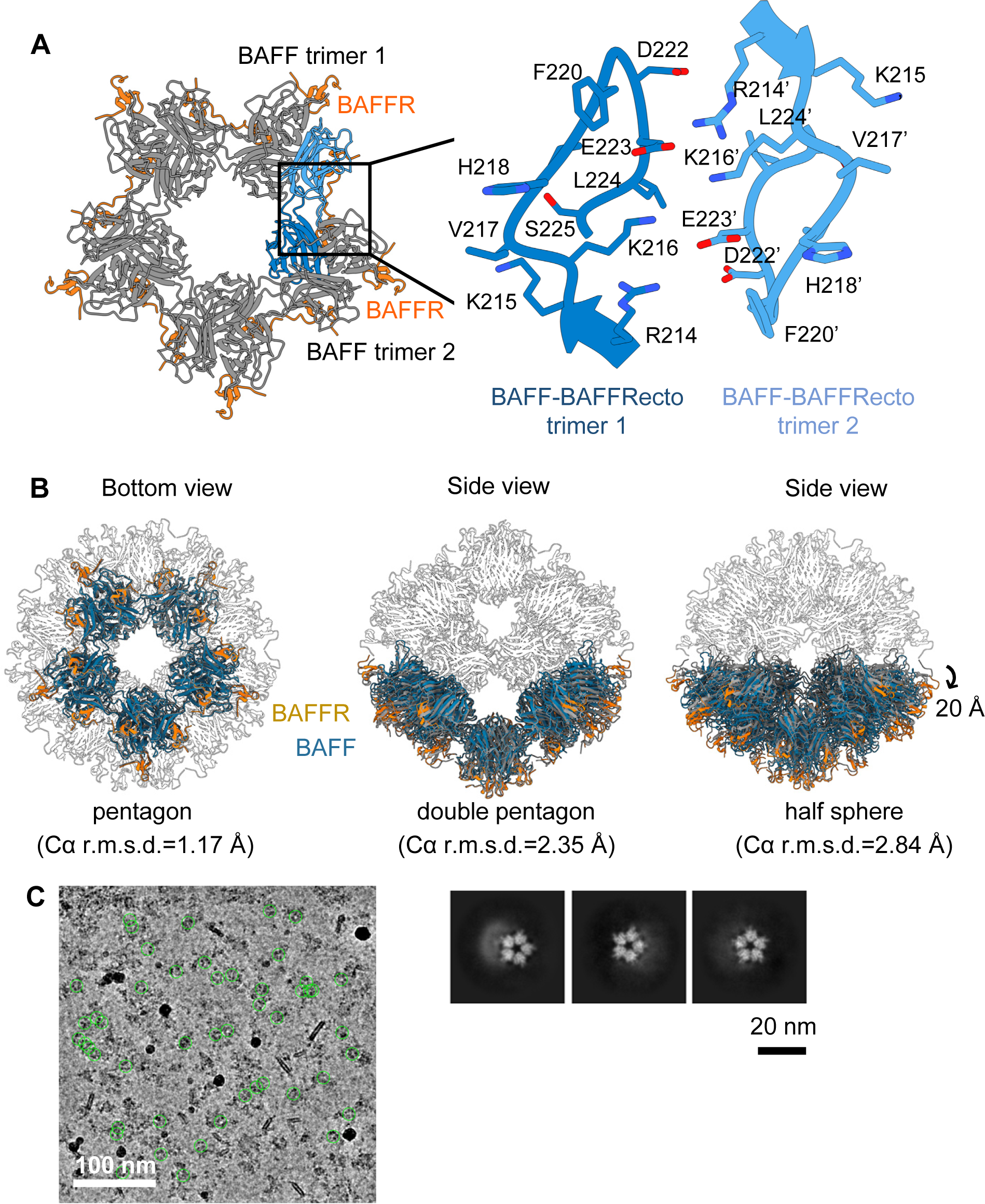
Structures of the BAFF-BAFFR clusters, related to Figure 4. (A) Close-up view of the BAFF clustering interface. Residue numbers for the second BAFF trimer are indicated with apostrophes. (B) Structural comparison between the pentagonal (left), double pentagonal (middle), and half-spherical (right) BAFF-BAFFRecto clusters and the full-size BAFF-BAFFRecto cage (PDB ID: 4V46). (C) A representative cryo-EM image (left) and 2D class averages (right three panels) of the BAFF-BAFFRfl complex. The protein particles selected for structure determination are indicated with green circles.

**Figure S5.**
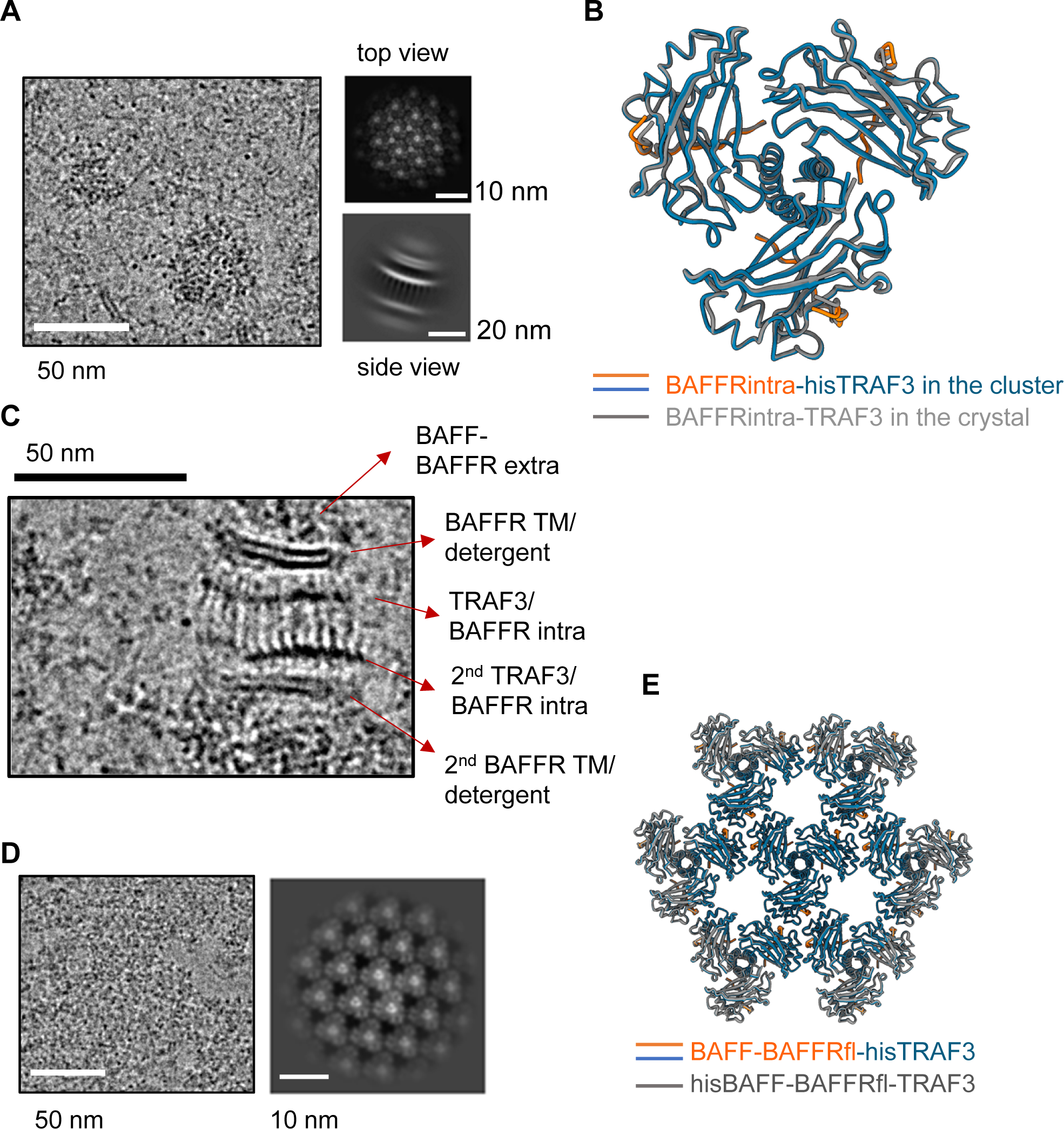
Structures of the hisBAFF-BAFFRfl-TRAF3 and BAFF-BAFFRfl-hisTRAF3 clusters, related to Figure 6. (A) A representative cryo-EM image and 2D class averages of the hisBAFF-BAFFRfl-TRAF3 complex. (B) Structural comparison between BAFFR-TRAF3 in the cluster and in the crystal structure (PDB ID: 2GKW). (C) A representative cryo-EM image showing the side view of the hisBAFF-BAFFRfl-TRAF3 cluster. (D) A representative cryo-EM image (left) and 2D class average (right) of the BAFF-BAFFRfl-hisTRAF3 complex. (E) Structural comparison of the BAFF-BAFFR-hisTRAF3 and hisBAFF-BAFFRfl-TRAF3 clusters.

## Methods S1: Cryo-EM processing schematics, related to Figures 1-7

**Processing Schematic 1.**
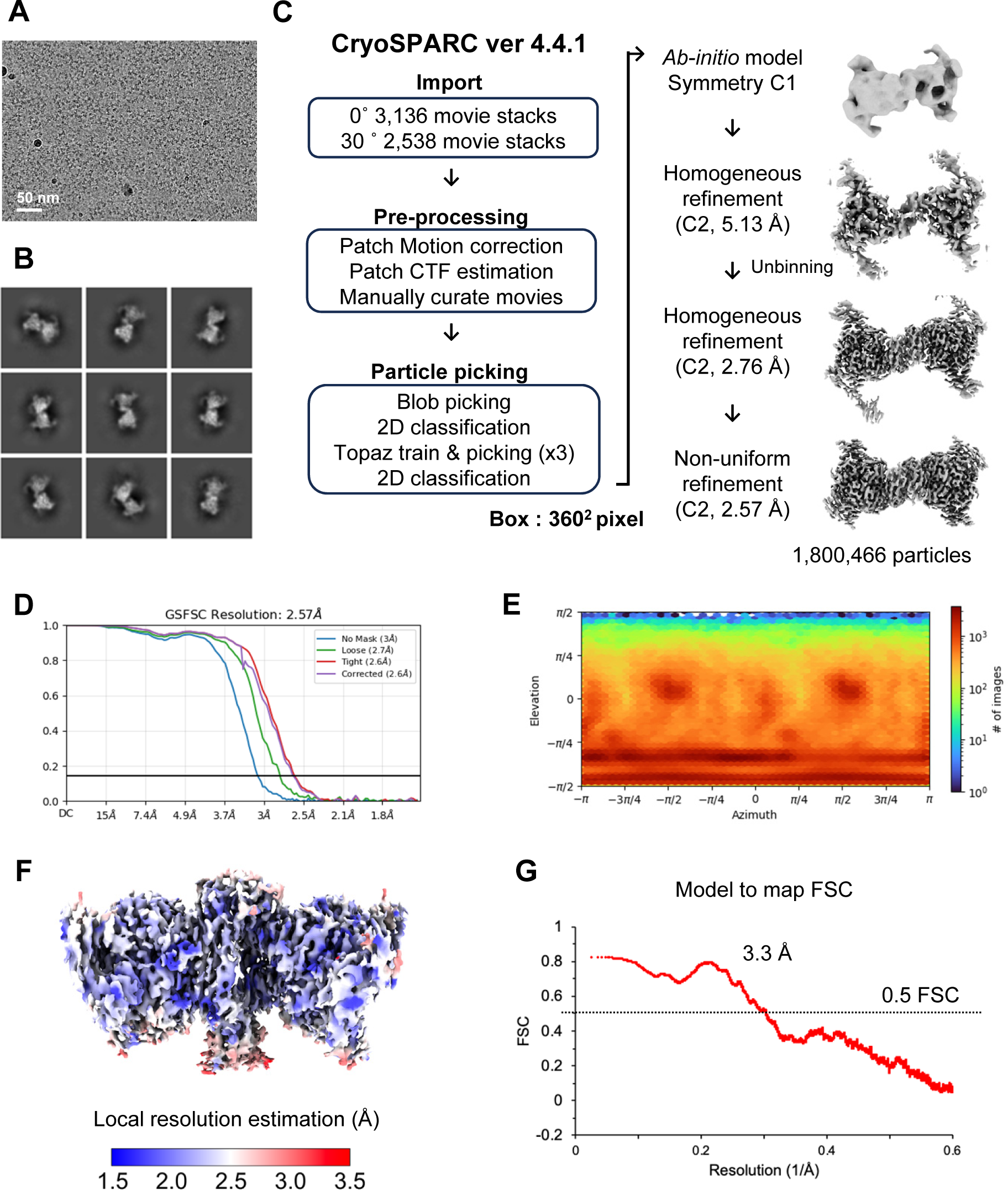
Cryo-EM data processing of the binary cluster of TNF*α*-TNFR1ecto complex. (A) A representative micrograph image (B) Selected 2D class averages (C) Summary of cryo-EM data processing (D) Map FSC curve (E) Orientation distribution of the particles used for 3D electron density reconstruction (F) Electron density map colored according to the local resolution estimation (G) The model to map FSC curve

**Processing Schematic 2.**
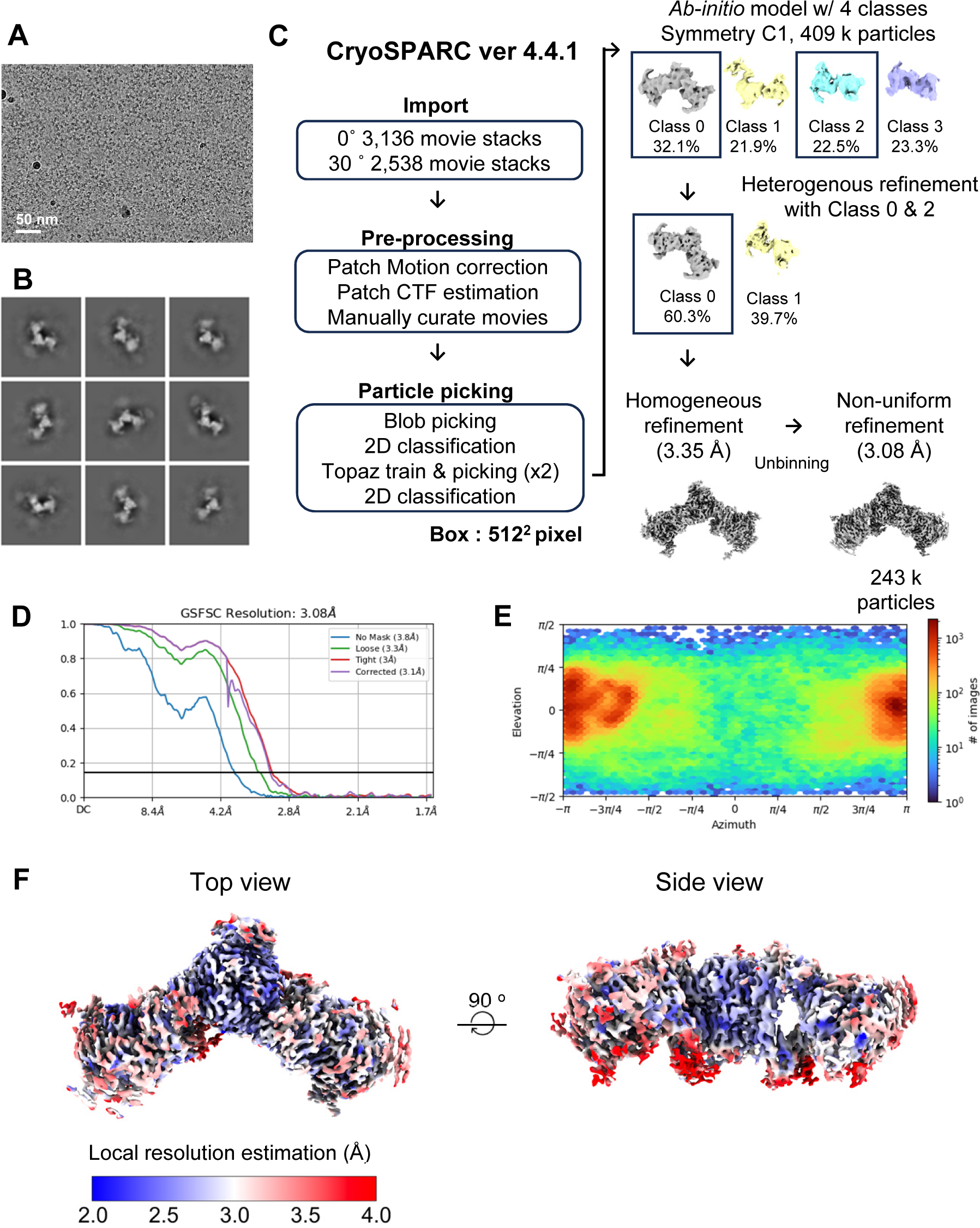
Cryo-EM data processing of the bent cluster of TNF*α*-TNFR1ecto. (A)A representative micrograph image (B) Selected 2D class averages (C) Summary of cryo-EM data processing (D) Map FSC curve (E) Orientation distribution of the particles used for 3D electron density reconstruction (F) Electron density map colored according to the local resolution estimation

**Processing Schematic 3.**
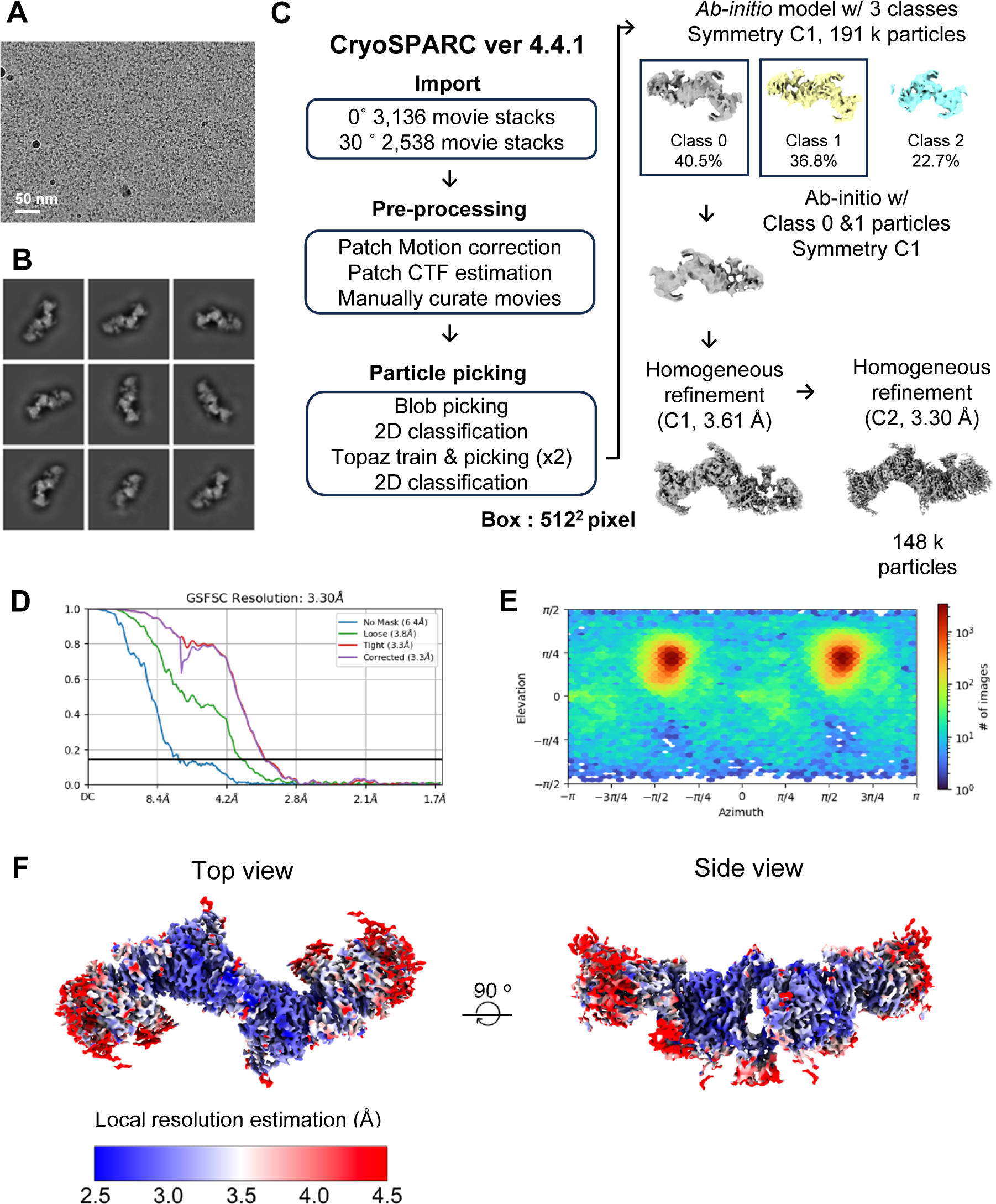
Cryo-EM data processing of the linear quadruple cluster of TNF*α*-TNFR1ecto. (A) A representative micrograph image (B) Selected 2D class averages (C) Summary of cryo-EM data processing (D) Map FSC curve (E) Orientation distribution of the particles used for 3D electron density reconstruction (F) Electron density map colored according to the local resolution estimation

**Processing Schematic 4.**
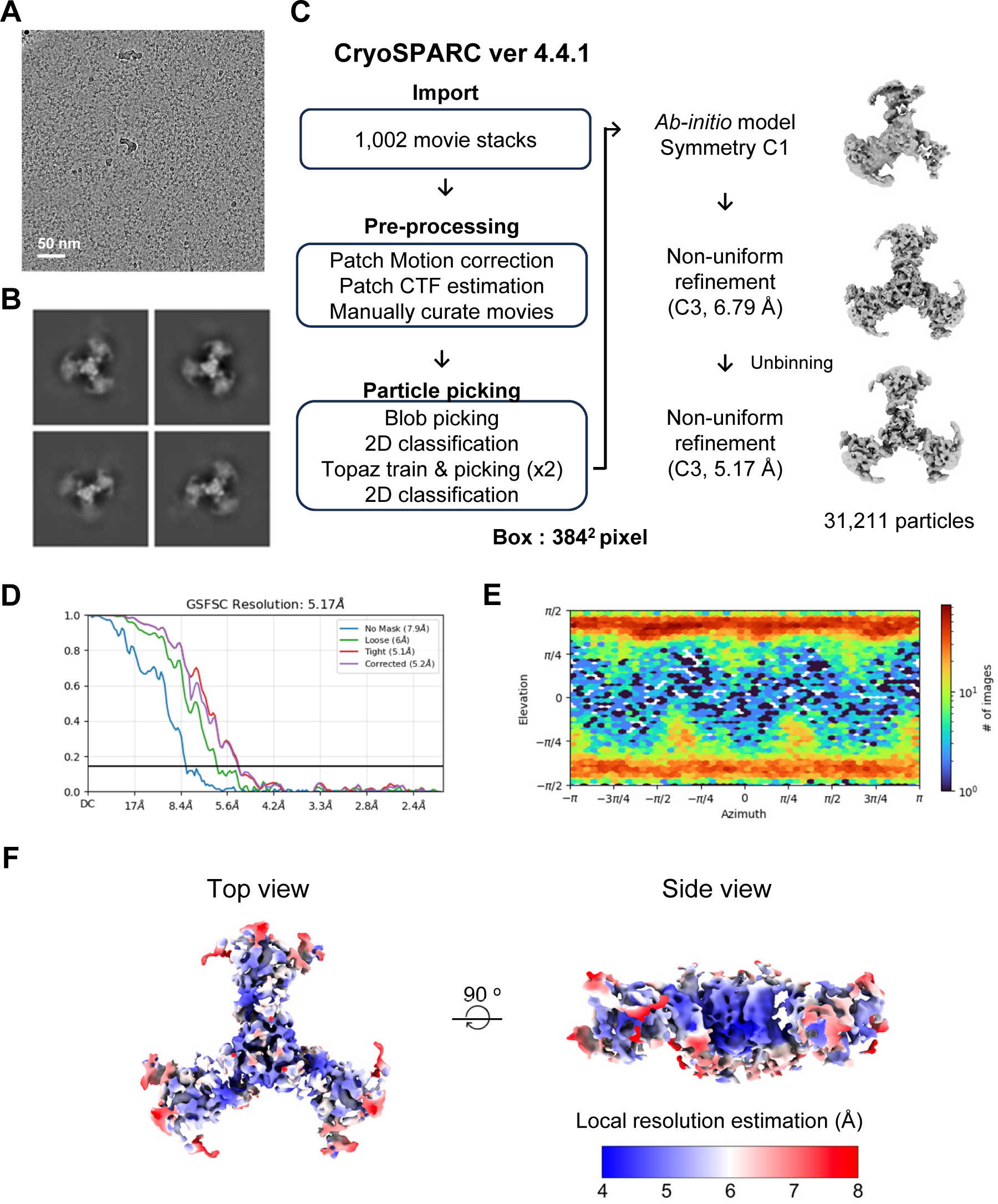
Cryo-EM data processing of the trigonal cluster of TNF*α*-TNFR1ecto. (A) A representative micrograph image (B) Selected 2D class averages (C) Summary of cryo-EM data processing (D) Map FSC curve (E) Orientation distribution of the particles used for 3D electron density reconstruction (F) Electron density map colored according to the local resolution estimation

**Processing Schematic 5.**
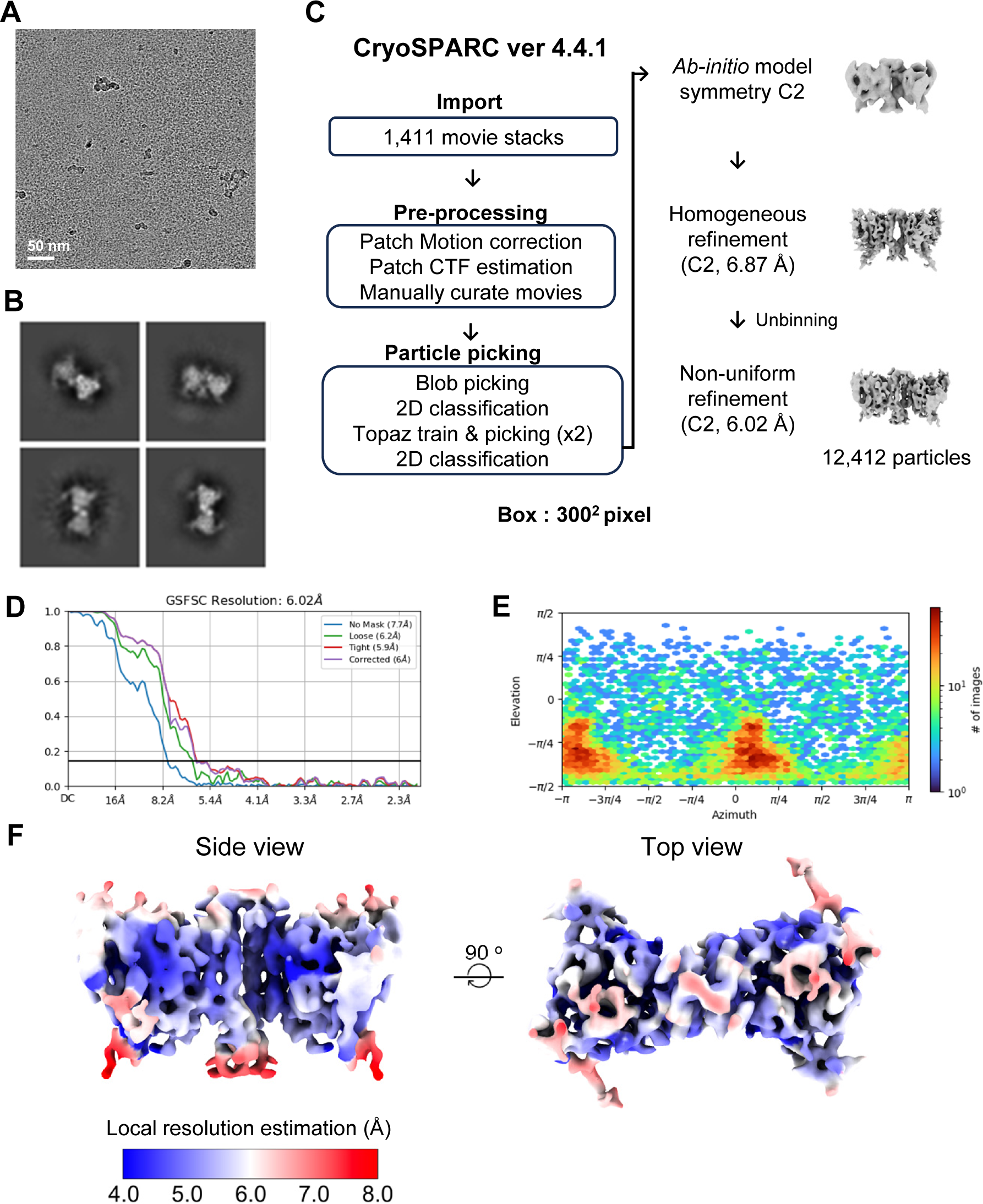
Cryo-EM data processing of the binary cluster of TNF*α*-TNFR1fl. (A) A representative micrograph image (B) Selected 2D class averages (C) Summary of cryo-EM data processing (D) Map FSC curve (E) Orientation distribution of the particles used for 3D electron density reconstruction (F) Electron density map colored according to the local resolution estimation

**Processing Schematic 6.**
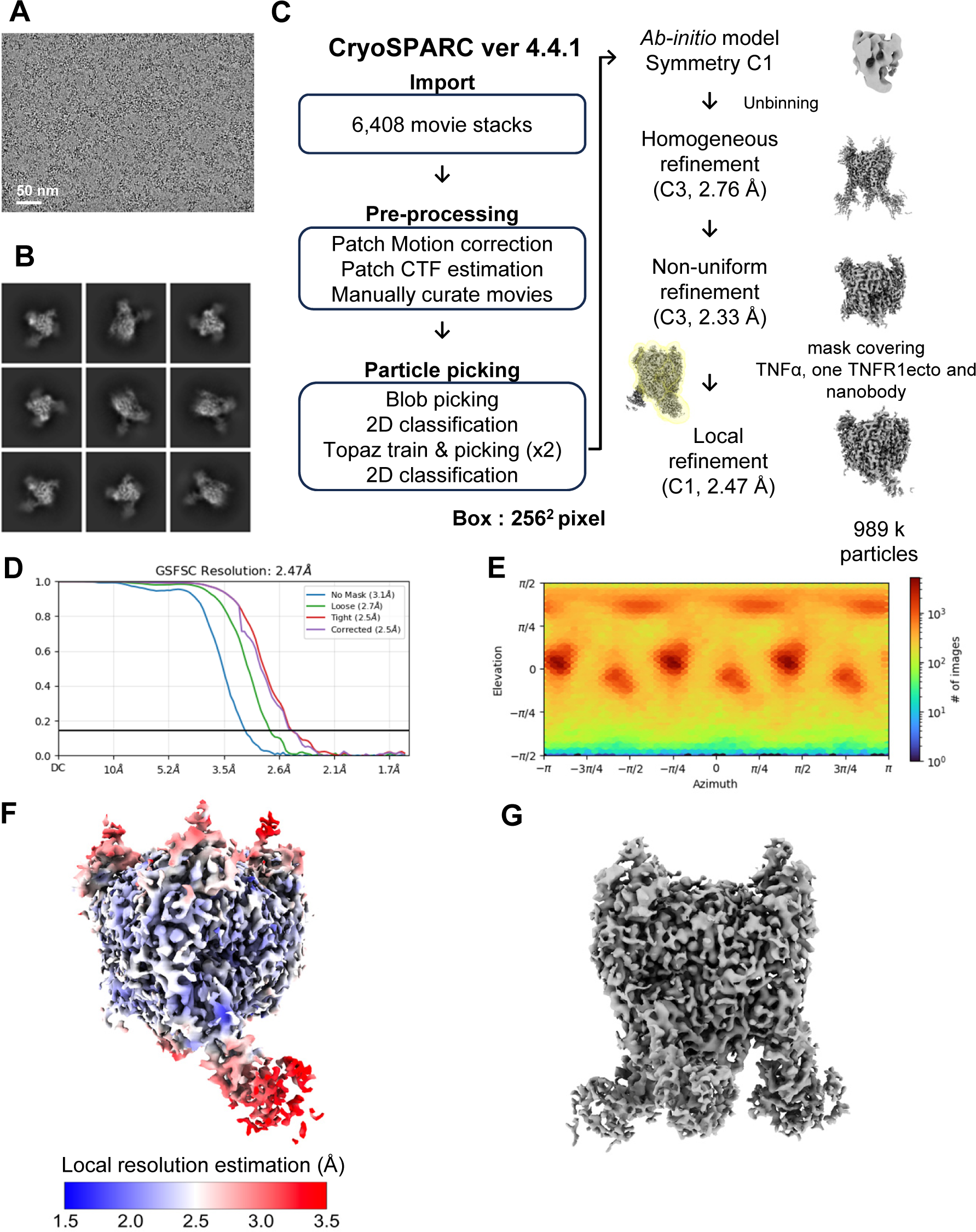
Cryo-EM data processing of the TNF-TNFR1ecto-DOM1h nanobody complex. (A) A representative micrograph image (B) Selected 2D class averages (C) Summary of cryo-EM data processing (D) Map FSC curve (E) Orientation distribution of the particles used for 3D electron density reconstruction (F) Electron density map colored according to the local resolution estimation (G) A composite map generated by imposing C3 symmetry

**Processing Schematic 7.**
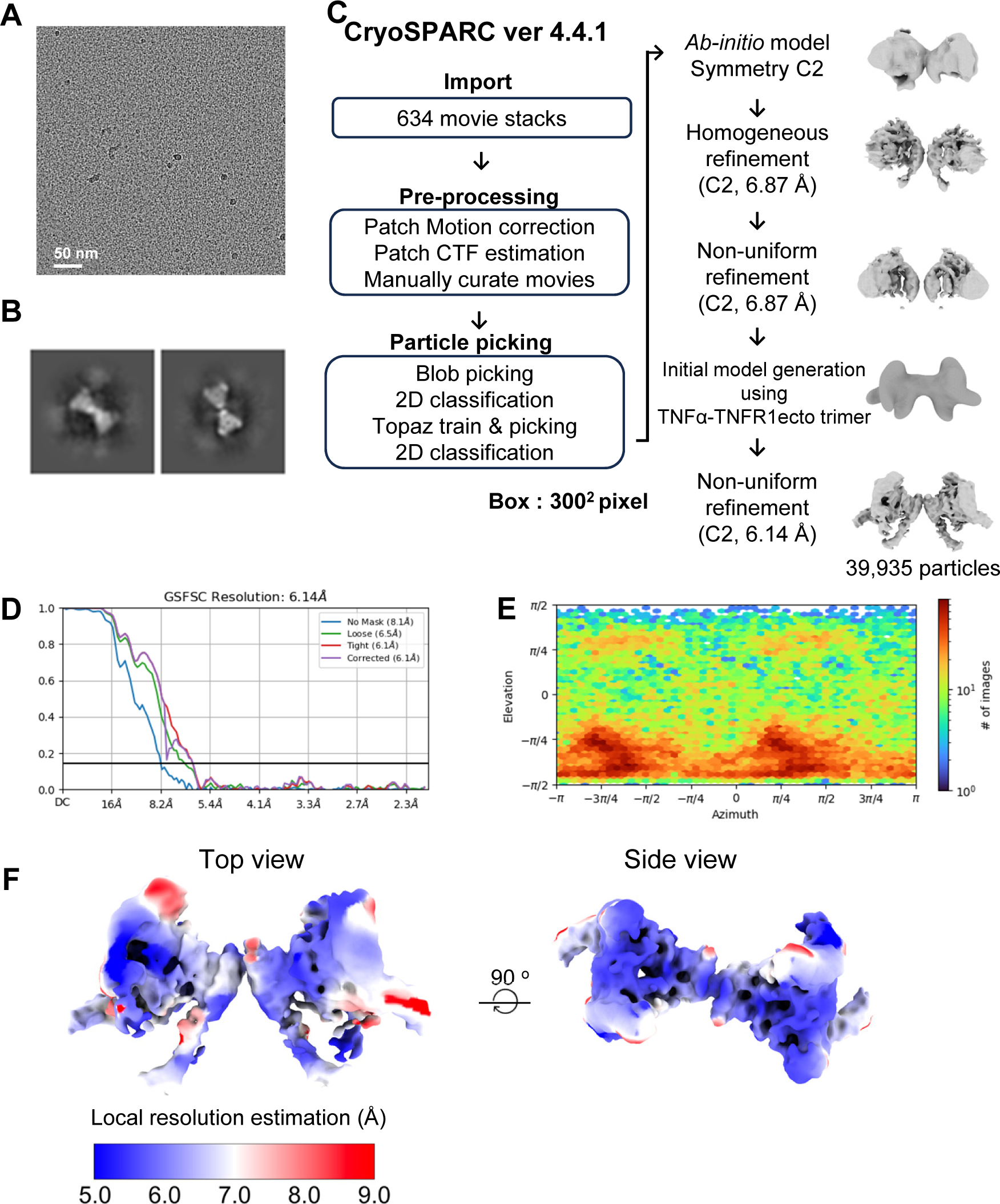
Cryo-EM data processing of the TNF*α*-TNFR1ecto-DOM1h nanobody complex on the lipid monolayer. (A) A representative micrograph image (B) Selected 2D class averages (C) Summary of cryo-EM data processing (D) Map FSC curve (E) Orientation distribution of the particles used for 3D electron density reconstruction (F) Electron density map colored according to the local resolution estimation

**Processing Schematic 8.**
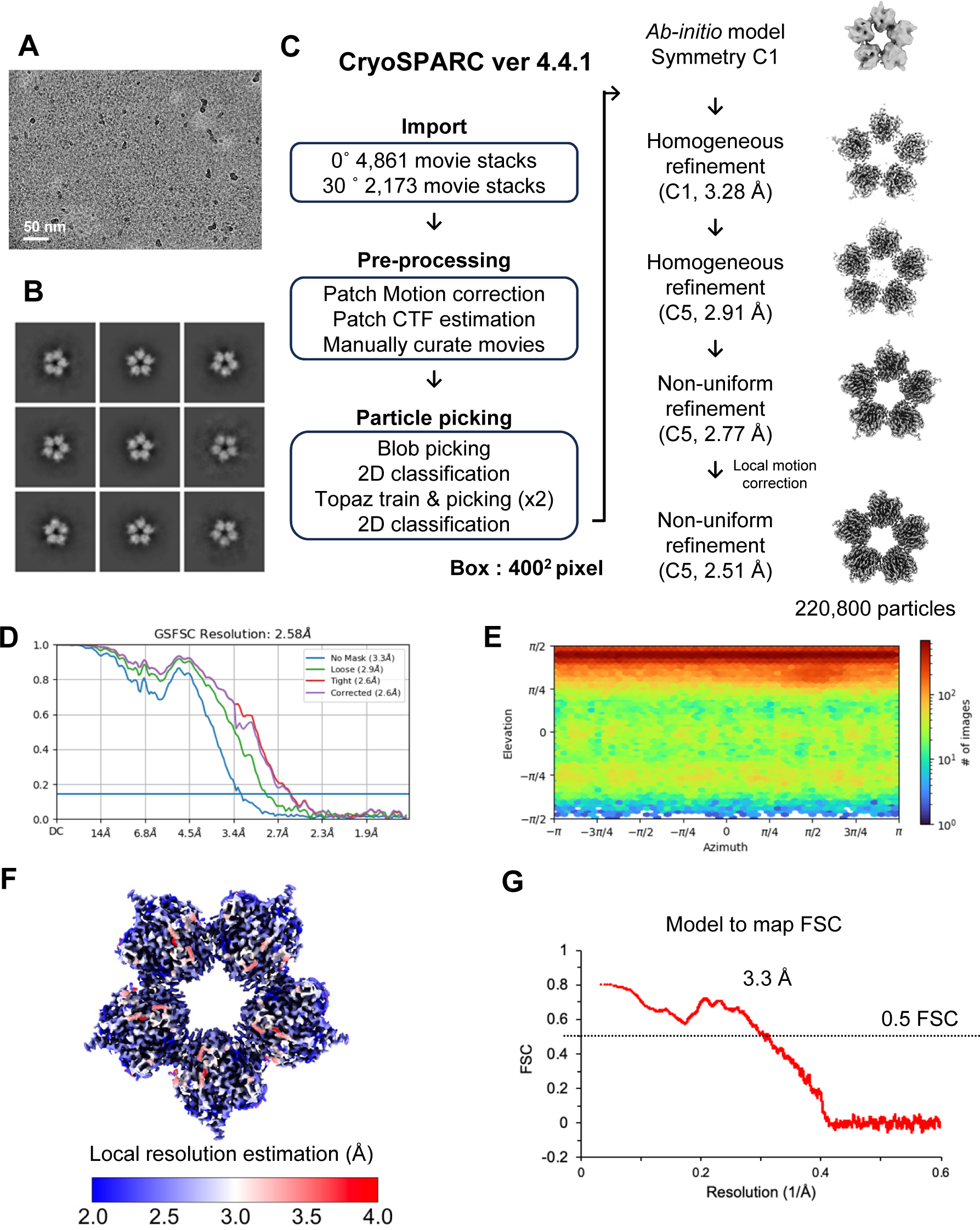
Cryo-EM data processing of the pentagonal cluster of BAFF-BAFFRecto. (A) A representative micrograph image (B) Selected 2D class averages (C) Summary of cryo-EM data processing (D) Map FSC curve (E) Orientation distribution of the particles used for 3D electron density reconstruction (F) Electron density map colored according to the local resolution estimation (G) The model to map FSC curve

**Processing Schematic 9.**
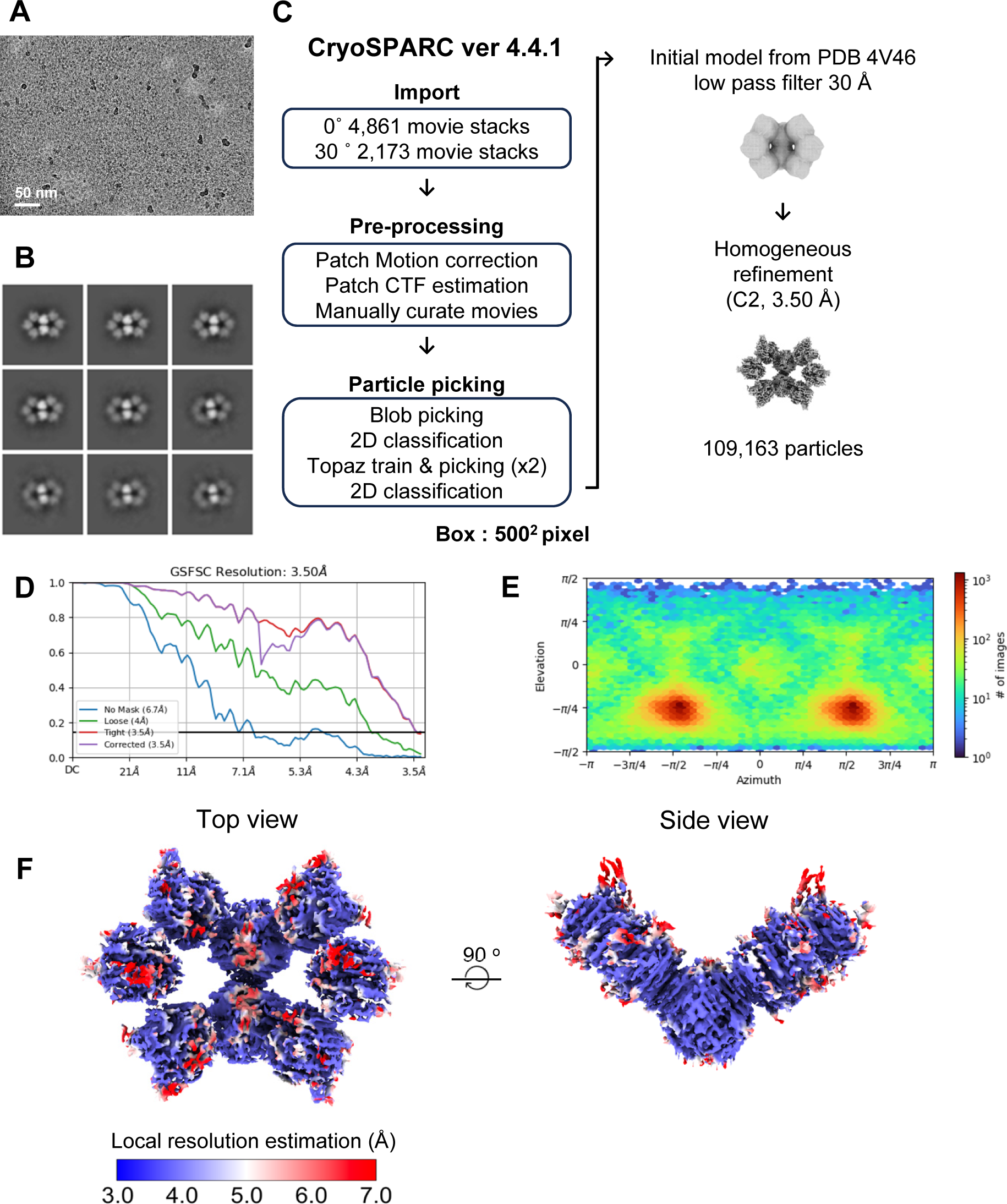
Cryo-EM data processing of the double pentagonal cluster of BAFF-BAFFRecto. (A) A representative micrograph image (B) Selected 2D class averages (C) Summary of cryo-EM data processing (D) Map FSC curve (E) Orientation distribution of the particles used for 3D electron density reconstruction (F) Electron density map colored according to the local resolution estimation

**Processing Schematic 10.**
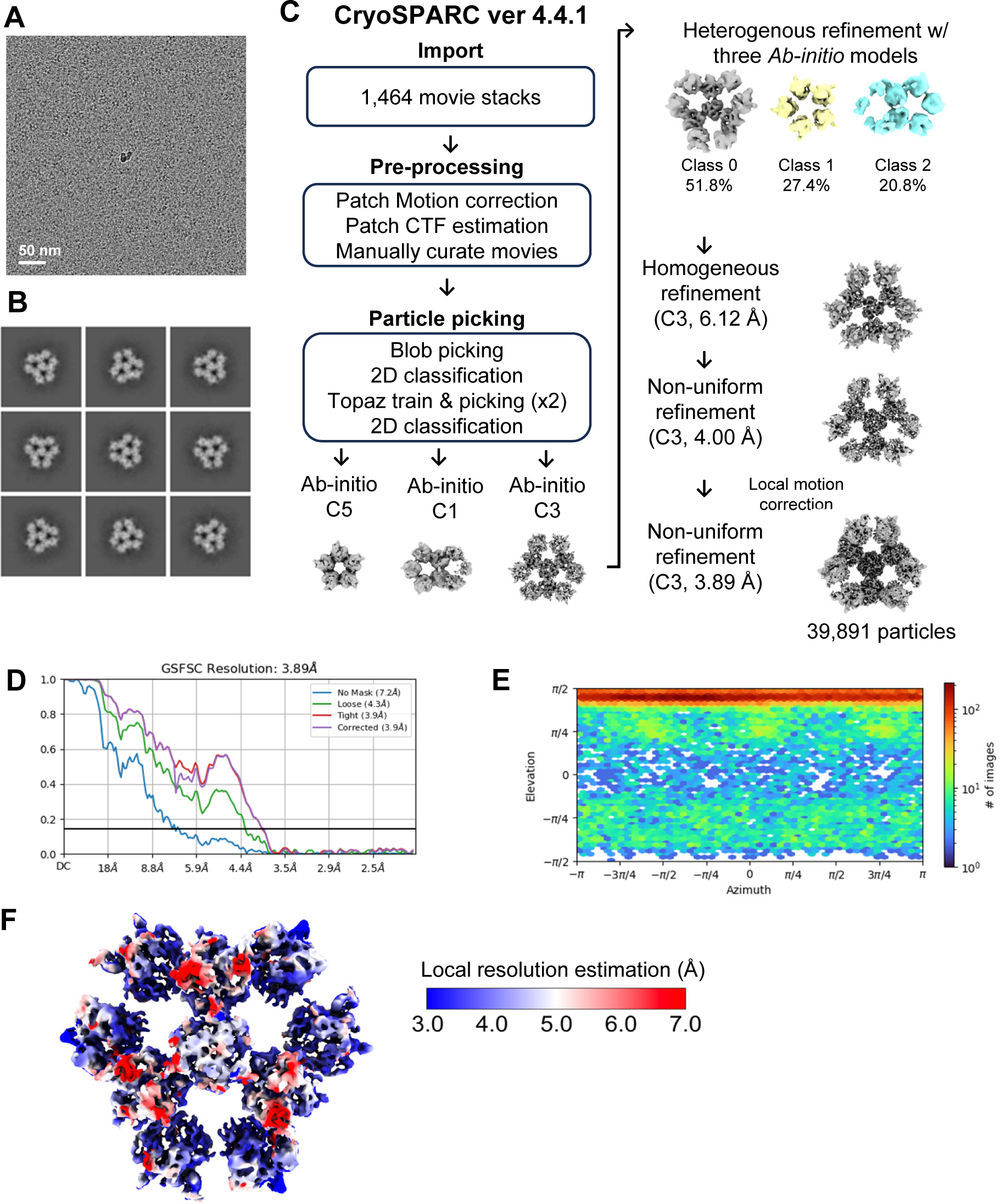
Cryo-EM data processing of the half sphere cluster of BAFF-BAFFRecto. (A) A representative micrograph image (B) Selected 2D class averages (C) Summary of cryo-EM data processing (D) Map FSC curve (E) Orientation distribution of the particles used for 3D electron density reconstruction (F) Electron density map colored according to the local resolution estimation

**Processing Schematic 11.**
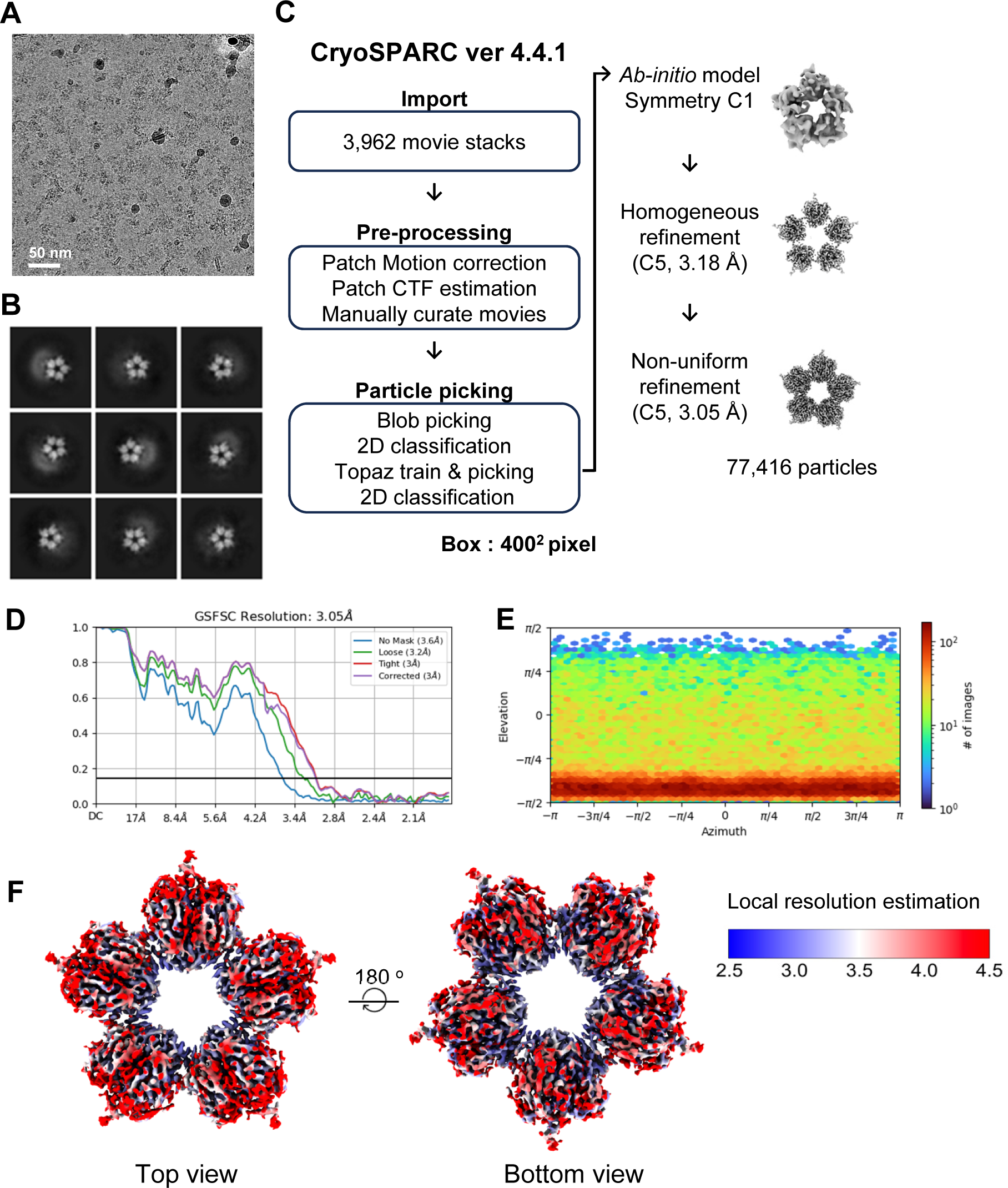
Cryo-EM data processing of the pentagonal cluster of BAFF-BAFFRfl. (A) A representative micrograph image (B) Selected 2D class averages (C) Summary of cryo-EM data processing (D) Map FSC curve (E) Orientation distribution of the particles used for 3D electron density reconstruction (F) Electron density map colored according to the local resolution estimation

**Processing Schematic 12.**
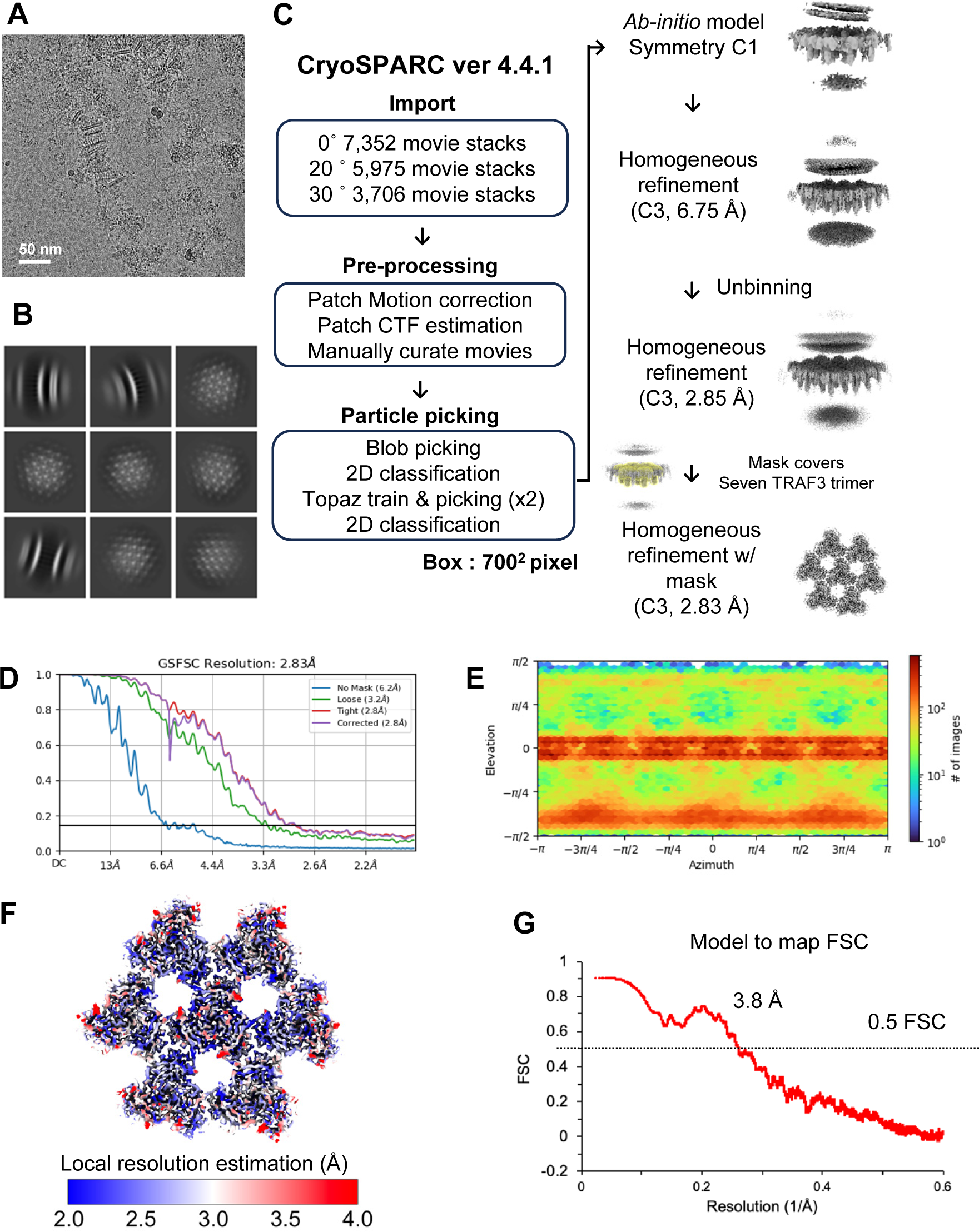
Cryo-EM data processing of the hisBAFF-BAFFRfl-TRAF3 cluster. (A) A representative micrograph image (B) Selected 2D class averages (C) Summary of cryo-EM data processing (D) Map FSC curve (E) Orientation distribution of the particles used for 3D electron density reconstruction (F) Electron density map colored according to the local resolution estimation (G) The model to map FSC curve

**Processing Schematic 13.**
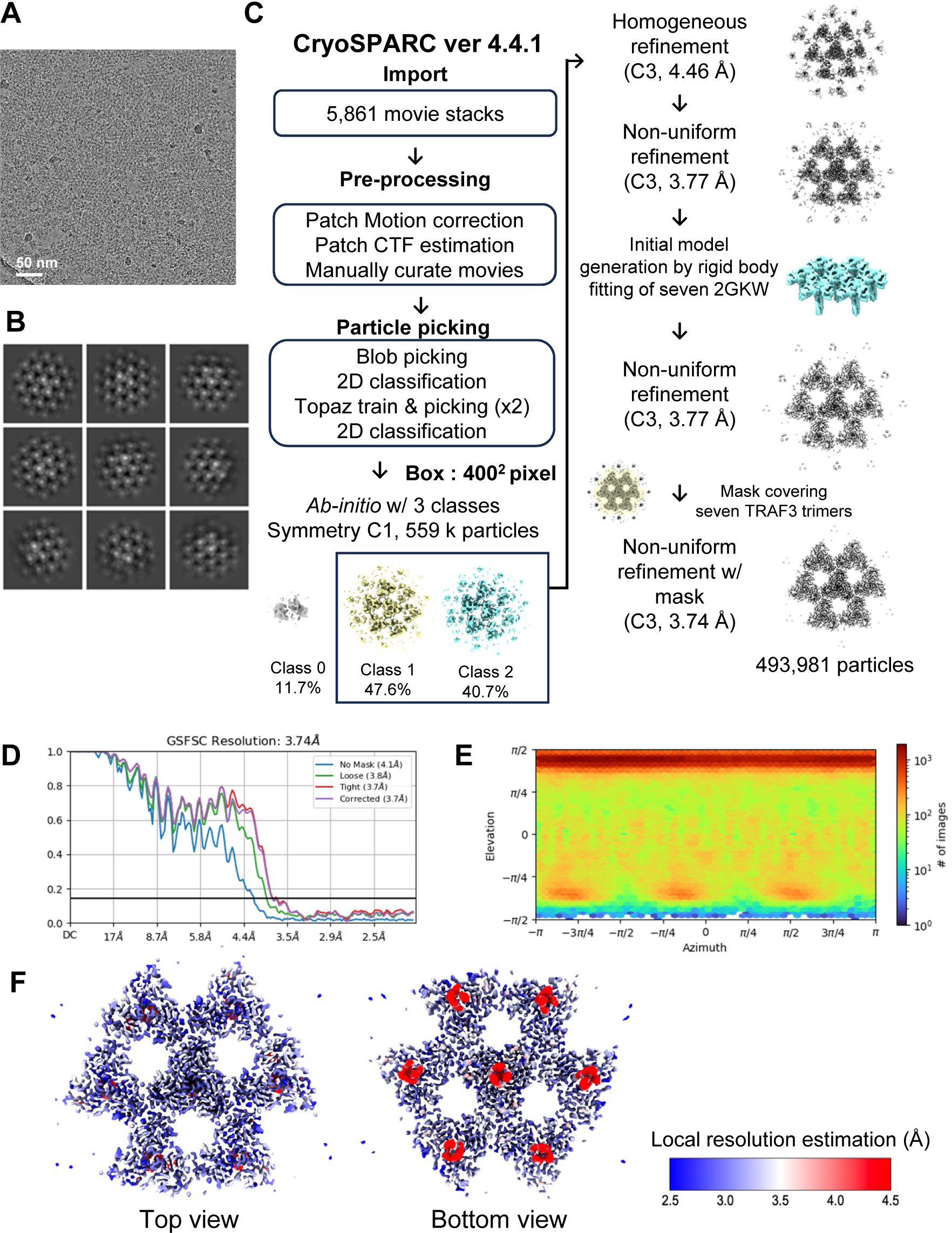
Cryo-EM data processing of the BAFF-BAFFRfl-hisTRAF3 cluster. (A) A representative micrograph image (B) Selected 2D class averages (C) Summary of cryo-EM data processing (D) Map FSC curve (E) Orientation distribution of the particles used for 3D electron density reconstruction (F) Electron density map colored according to the local resolution estimation

